# Multimodal screen reveals noise regulatory proteins

**DOI:** 10.1101/2024.07.17.603871

**Authors:** Óscar García-Blay, Xinyu Hu, Christin L. Wassermann, Tom van Bokhoven, Fréderique M.B. Struijs, Maike M.K. Hansen

## Abstract

Gene-expression noise can influence cell-fate choices across pathology and physiology. However, a crucial question persists: do regulatory proteins or pathways exist that control noise independently of mean expression levels? Our integrative approach, combining single-cell RNA sequencing with proteomics and regulator enrichment analysis, reveals 32 putative noise regulators. SON, a nuclear speckle-associated protein, alters transcriptional noise without changing mean expression levels. Furthermore, SON’s noise regulation can propagate to the protein level. Long-read and total RNA sequencing shows that SON’s noise regulation does not significantly change isoform usage or splicing efficiency. Moreover, SON depletion reduces state-switching in pluripotent mouse embryonic stem cells and impacts their fate choice during differentiation. Collectively, we discover a class of proteins that regulates noise orthogonally to mean expression levels. This work serves as a proof-of-concept that can identify other functional noise-regulators throughout development and disease progression.

## INTRODUCTION

Fluctuations in gene expression––termed “noise”^1,2^––have been shown to play a critical role in a range of biological systems from microbes to higher eukaryotes.^3–7^ In eukaryotic developmental processes, cells utilize noise to populate functionally distinct states.^8–10^ Specifically, amplified noise potentiates and accelerates lineage specification^11^ while attenuated noise ensures stability of lineage commitment.^12^ Mounting evidence shows that gene-expression noise is a universal aspect of single-cell biology. The existence of mechanisms that alter gene expression noise,^13–20^ including feedback topologies that regulate locus-specific noise, suggest that gene-expression noise can be modulated by proteins.^5,21,22^ Yet, it remains unclear if specific proteins, or protein networks exist that globally regulate gene-expression noise (i.e., independent of the mean).

Here, we develop a screening strategy to identify noise-regulatory proteins in mouse embryonic stem cells (mESCs). The approach utilizes global translation inhibition^23^ (i.e., perturbation of potential protein regulators), and single-cell RNA sequencing (scRNA-seq) to quantify the changes in noise of all transcripts (i.e., potential mRNA targets). Remarkably, the screen identifies a subset of transcripts whose noise is regulated by proteins in a mean-independent manner. Subsequent proteomics and regulator enrichment analysis classify 32 noise-regulators spanning diverse functions including DNA methylation, nuclear compartmentalization and splicing. The most highly-ranked noise regulator, SON, is a protein that has been linked to nuclear speckle assembly^24,25^ and splicing regulation^26,27^. SON depletion appears to impact developmental cell-fate choices with implications in mESC pluripotency and differentiation. Taken together, this study reveals a class of proteins that regulate gene expression noise orthogonally to mean expression levels. The discovery of noise regulators implies that proteins may have been selected for their ability to regulate gene expression noise across health and disease.^7,28–30^

## RESULTS

### Blocking translation reveals a set of transcripts whose noise is regulated by proteins

To determine how noise regulators might be identified, we hypothesized that noise modulation can be categorized into two distinct types: noise enhancement and noise repression (**Figure 1A**, top). Noise enhancement amplifies fluctuations in gene expression levels, providing cells with a greater degree of exploratory potential (**Figure 1A**, blue arrow), which increases the probability of a cell surpassing a phenotypic threshold (**Figure 1A**, orange dashed line) and generates higher phenotypic diversity.^11,31^ Conversely, noise repression (**Figure 1A**, bottom) reduces the amplitude of these fluctuations, consequently limiting the exploratory potential and generating a phenotypically more homogeneous distribution of cells.^21,31^ If a protein is a noise regulator (either enhancer or repressor) that acts on a target mRNA, we expect that removing translation of the protein regulator (**Figure 1A**, middle, yellow line) abolishes the noise regulation effect on its target mRNA (**Figure 1A**, middle).^32^ As a result, the noise levels of the target mRNA should change, while mean expression levels remain unchanged (**Figure 1A**, middle compared to right). Therefore, we define a noise regulator as a protein that modulates noise levels of a target mRNA without altering mean levels of that same target. Importantly, we do not exclude that a regulator can act as both a mean and a noise regulator of two different target mRNAs.

**Figure 1.**
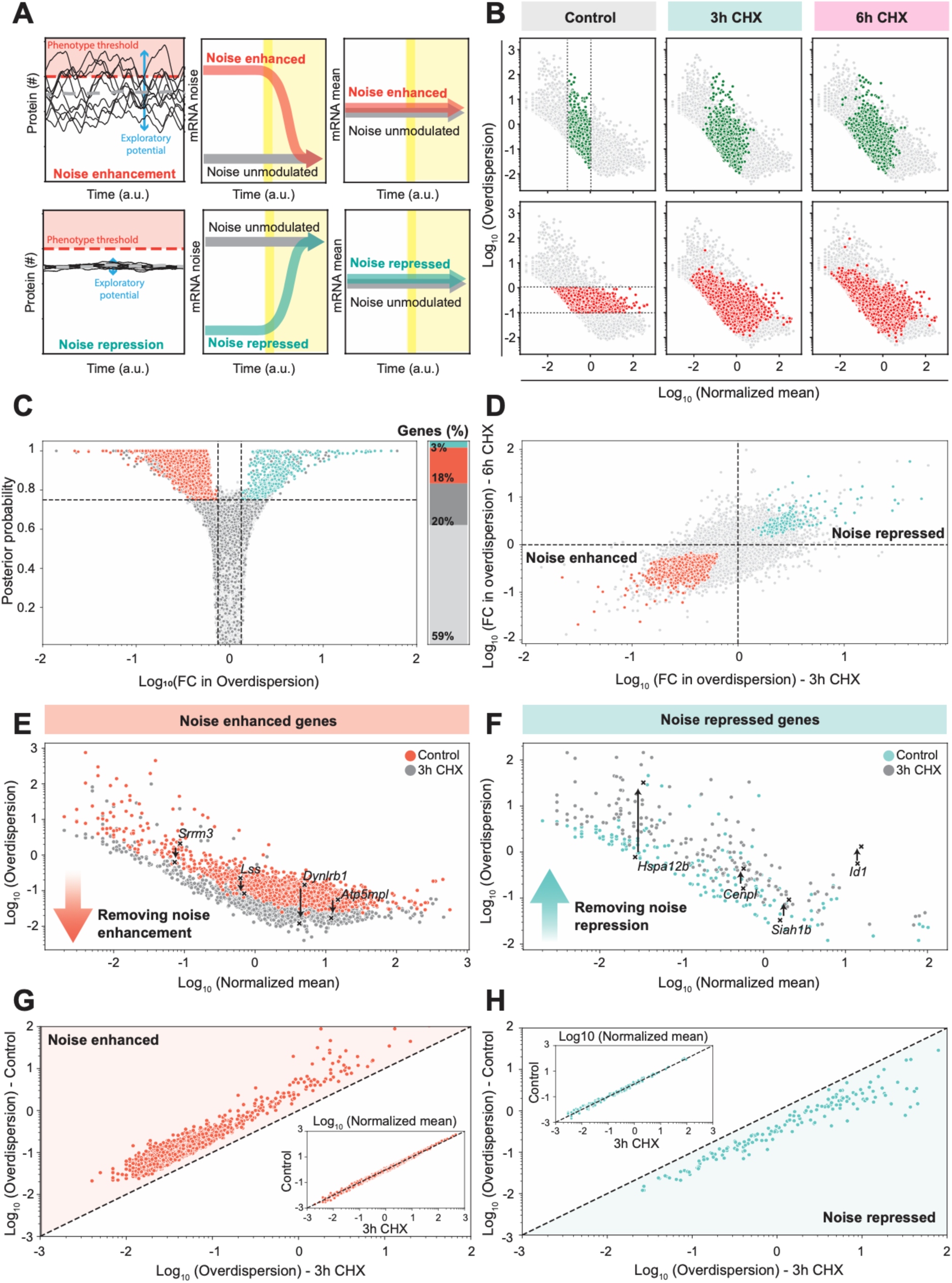
Blocking translation reveals transcripts whose noise is controlled by proteins. **(A)** Schematic illustrating noise enhancement (left, top) and noise repression (left, bottom). Black lines represent single-cell protein expression levels and the population mean is denoted by the grey dashed line. Larger fluctuations increase the exploratory potential of a cell and drive cell-state transitions across a phenotypic threshold (orange dashed line). When noise regulation is perturbed by blocking translation (yellow shaded area) noise will change in the opposite direction of the noise regulation mechanism (center), while mean expression levels remain constant (right). **(B)** Normalized mean expression versus overdispersion from scRNA-seq of mESCs from control sample and treated with 250 µM of CHX for 3 and 6 hours (1462, 1560, 2395 cells respectively) with highlighted genes displaying a central range in mean expression (top, green) or overdispersion (bottom, red) in the control. **(C)** Differential expression analysis with BASiCs of the 3-hour CHX treated sample compared to the control. Volcano plot represents the fold change in overdispersion versus the certainty (Posterior probability) that this change is significant (dashed lines are the significance thresholds). Stacked bar plot represents the % genes that display significant changes in overdispersion (orange: 2935 noise-enhanced genes; turquoise: 522 noise-repressed genes; dark grey: 3398 genes excluded because they display differential mean expression). **(D)** Fold change in overdispersion of the 3-hour CHX sample compared to control versus the 6-hour CHX sample compared to the control. Genes showing a consistent significant change in overdispersion after both 3 and 6 hours of CHX treatment are highlighted in turquoise (166 noise-repressed genes) and orange (1484 noise-enhanced genes). **(E-H)** scRNA-seq of mESCs from control sample and treated with 250 µM CHX (turquoise or orange) for 3 hours. Mean expression versus noise (overdispersion) of 1484 genes classified as noise-enhanced (E) and 166 genes classified as noise-repressed (F). These genes change in noise levels (G-H) without changing in mean mRNA expression levels (G-H, inset).

To identify which target mRNAs might be noise-enhanced or -repressed in mESCs, we treated cells with the translation inhibitor Cycloheximide^23,33^ (CHX, 250 µM) and performed scRNA-seq.^34^ Since CHX is dissolved in ethanol (EtOH) the control samples were also exposed to EtOH. Translation is completely blocked within 30 minutes of treatment, as demonstrated by the lack of incorporation of the amino acid analogue homopropargylglycine (HPG) (**Figure S1A**). CHX quickly induces the decay of short-lived proteins such as NANOG (**Figure S1B**). Importantly, 3-6 hours of CHX treatment has a minimal effect on cell viability (**Figure S1C**). Fluorescent staining of newly transcribed RNA via metabolic labelling with 5-ethynyluridine (5EU) reveals no change in global RNA degradation in the presence of CHX (250 µM) compared to the control (**Figure S1D-E**). While global RNA synthesis is slightly diminished upon CHX treatment (**Figure S1F-G**), mRNA transcription appears mostly unperturbed, in particular when compared to the transcriptional inhibitor Actinomycin D (ACT-D)^35^ (**Figure S1H-J**), consistent with previous findings.^36^ Although there are no changes in global mRNA abundance, we cannot exclude that CHX treatment increases mRNA abundance for some genes while decreasing it for others as a consequence of changes in transcript specific stability upon translation inhibition.

Having established that short (3-6 hour) CHX treatment blocks translation, decreases protein abundance while maintaining mRNA transcriptional activity and cell viability, we next performed scRNA-seq after blocking translation for 3 and 6 hours (**Figure S2A**). After UMAP dimensionality reduction of the data, the CHX treated samples clustered separately from the control in both bulk RNA-seq (**Figure S2B**) as well as scRNA-seq experiments (**Figure S2C**). This suggests that overall changes in gene expression occur progressively from 3 to 6 hours of CHX treatment. When analyzing single-cell gene expression levels, both mean mRNA expression and noise (quantified as overdispersion) change over time (**Figure 1B and Figure S2D**). Overdispersion is quantified as the excess noise not explained by the reference model for technical noise^37^ and behaves similarly to CV^2^ (=σ^2^/μ^2^). Low expressed genes tend to increase in mean mRNA abundance upon CHX treatment, indicating that they are being actively repressed in untreated conditions, while the inverse appears to be true for highly expressed genes^38^ (**Figure 1B**, top). Conversely, genes that are moderate in their expression levels either increase or decrease in mean mRNA abundance when treated with CHX (**Figure S2D**, top), indicating more diverse mean-regulation mechanisms. Interestingly, the same holds for noise-regulated genes (**Figure 1B** and **Figure S2D**, bottom). Notably, changes in mean mRNA expression occur progressively (**Figure S2E**, top compared to bottom) and are consistent with bulk RNA-seq results (**Figure S2F**) as well as over time (**Figure S2G**). Conversely, most changes in mRNA noise take place within the first 3 hours of treatment (**Figure 1C** compared to **Figure S2H**). Together, these results indicate that blocking translation for 3-6 hours is sufficient to reproducibly identify relatively fast (i.e., 3-hour) changes in mRNA levels.

To identify transcripts that are solely noise-modulated after removing technical noise^37^, genes that show significant changes in mean for both the 3- and 6-hour CHX treatment (20% and 35% of the genes respectively) were excluded from the analysis (**Figure S2E**). Of the remaining genes that display no significant change in mean mRNA expression (80% and 65%, for 3- and 6-hour CHX treatment respectively), 15-18% of transcripts present significant changes in noise with 3 and 6 hours of treatment (**Figure 1C** and **Figure S2H**). In total, 1650 genes (∼10%) show consistent significant changes in noise after both 3 and 6 hours of CHX treatment (**Figure 1D**). This set of noise-regulated transcripts could be divided into two groups:

1. *inherently high noise* transcripts that are *noise-repressed* in untreated conditions and increase back to their basal (i.e., unregulated) noise levels when translation is blocked (**Figure 1E**); and
2. *inherently low noise genes* that are *noise-enhanced* in untreated conditions and decrease back to their basal noise levels when translation is blocked (**Figure 1F**). Both groups of transcripts display a clear change in noise (**Figure 1G-H**), while mean mRNA levels remain comparable to the control (**Figure 1G-H**, inset). Interestingly, the group of noise-enhanced transcripts (1484 genes, ∼9%) is much more prominent than the group of noise-repressed transcripts (166 genes, ∼1%). The finding that noise enhancement appears ∼9-fold more abundant than noise repression, could be explained by the latter being more difficult to mechanistically decouple from mean modulation or by noise repression requiring longer perturbations. Moreover, noise enhancing mechanisms could simply be more prevalent in mESCs, as previous literature has described high levels of noise as biologically relevant in these cells,^8,39–41^ and noise reduction might become more important during lineage commitment.^12,40^ Collectively, these data show that a previously underappreciated portion of the transcriptome appear to be noise-regulated.

### Transcriptome-wide screen uncovers 32 noise regulators

After classifying this set of transcripts as noise-regulated, we sought to determine the proteins responsible for the observed noise changes. We thus performed two consecutive analyses: i) determined which proteins quickly decrease in abundance within 3-6 hours of CHX treatment (i.e., are short lived); and ii) identified which regulators are enriched with noise-regulated mRNA targets. First, we performed proteomics after translation inhibition for 3-9 hours and detected 5,323 proteins in all 4 treatments (4 replicates per treatment) (**Figure S3A**). We found ∼1,000 proteins that show a significant relative depletion after CHX treatment (**Figure 2A** and **Figure S3B**), consistent with previous findings.^33^ These ∼1,000 *depleted proteins* were labeled as *possible protein regulators.* Next, we performed a regulator enrichment analysis for the separate *target lists* (established through scRNA-seq), (**Figure 2B**, see STAR Methods for more details). In short, for each *possible protein regulator* the *known target list* was obtained from existing datasets.^42–45^ This list of known targets was compared to the list of *noise-enhanced mRNA targets* as well as multiple *control lists* consisting of *unchanged targets*. Enrichment was computed and a protein was classified as a noise regulator if it showed an Odds ratio > 1, with a false discovery rate (FDR) <0.05 and the probability of classifying that regulator as a noise regulator from 10 randomized control lists was ≤0.1.

**Figure 2.**
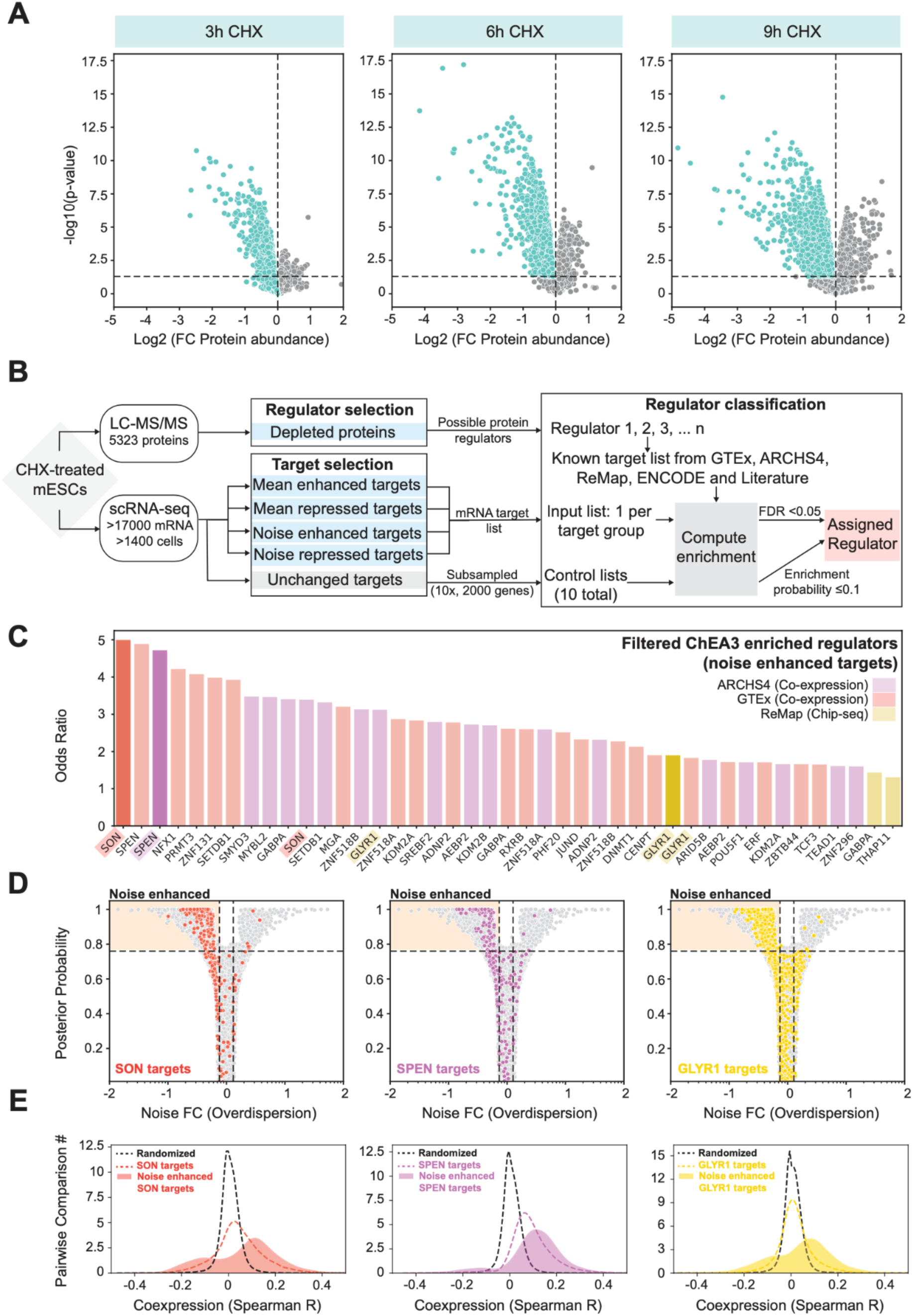
Proteomics and regulator enrichment analysis uncovers proteins that regulate noise. (A) Fold change in protein abundance versus p-value for mESCs treated with 250 µM of CHX for 3 (left), 6 (middle), and 9 (right) hours compared to the control (dashed lines are the significance thresholds), of 5323 proteins detected in LC-MS/MS analysis. (B) Schematic overview of analysis performed to identify noise regulators. (C) Noise regulators identified through regulator enrichment analysis ranked according to their odds ratio. The color represents the database from which the regulator and respective target set was obtained. (D) Fold change in overdispersion and certainty (Posterior probability) of 3h CHX sample compared to the control. Highlighted genes are targets of SON (orange; left), SPEN (violet; middle) or GLYR1 (yellow; right) classified by GTEx, ARCHS4, or ReMap respectively. (E) Spearman correlation values from pairwise comparisons of target mRNA expression levels of SON (dashed orange line; left), SPEN (dashed violet line; middle) or GLYR1 (dashed yellow line; right), classified by GTEx, ARCHS4, or ReMap respectively. Full colored distribution represents the subset of targets that are defined as noise-enhanced. Black dashed distributions represent randomized set of genes.

This analysis yielded a list of 32 potential noise-regulatory proteins (**Figure 2C**) from three different databases (ARCHS4, GTEx, and ReMap) included in ChEA3.^42–45^ When performing a similar analysis for the mean-regulated transcripts we identified known transcriptional activators in mESCs, confirming the validity of this approach (**Figure S3C**; **Table S1**). Interestingly, the list of noise regulators spans transcription factors (i.e., POU5F1/OCT4^46^, TCF3^47^), methylation associated proteins (i.e., SETDB1^48^, SMYD3^49^, GLYR1^50^), and proteins involved in splicing (SON^26,27^) or nuclear speckles/compartments (SON^25^, SPEN^51^). Analysis of targets shared between the identified noise regulators revealed that while some regulators present a high target-set overlap with other regulators (e.g., KDM2A in GTEX database or MYBL2 in ARCHS4 database), others show a comparable overlap as is expected from randomized controls (e.g., SON in GTEX database) (**Figure S3D-E**). This indicates that noise modulation might be enforced through synergistic, antagonistic or redundant regulators, but also occurs in isolation. Notably, a single regulator does not necessarily alter only one kinetic rate, allowing for the possibility of noise modulation through multiple kinetic steps but involving a single regulator.^11^ Lastly, while mean regulators show enrichment in Gene Ontology terms related to regulation of transcription or known processes in development, noise modulators show no significant enrichment of a particular function (**Figure S3F**, orange; **Table S1**), indicating a possible underexplored area of biological regulation.

The three most highly ranked protein regulators of each database––SON, SPEN and GLYR1– –show fast decay upon CHX treatment (**Figure S3G**), as well as an enrichment of mRNA targets in the transcript set that is noise-enhanced (**Figure 2D**). Furthermore, both their full set of known targets (defined by the corresponding database) and the subset of noise-regulated targets (**Figure 2D**, shaded area) show stronger co-expression with respect to a randomized control in the untreated scRNA-seq sample (**Figure 2E**). This indicates a functional relationship behind the expression patterns of these genes through a common regulator, in this case SON, SPEN or GLYR1 respectively.^52,53^ Together these data highlight that this screening approach can be implemented to classify proteins as noise regulators––regardless of their mode of action––as long as regulator-target relationships have previously been identified.

### SON modulates noise without affecting mean mRNA and protein expression

Having established a list of noise-regulators, we selected the most highly ranked protein SON, for more extensive analysis. In order to study the noise modulation functions of SON, we performed an siRNA mediated knock down (KD)^54^ for 24 hours (**Figure 3A**). scRNA-seq of both KD and control cells was subsequently performed and as expected *Son* KD is also observable at the RNA level (**Figure 3B**). Noticeably, the mean mRNA expression after 24 hours of SON depletion does not change considerably (**Figure 3C**, top). The differential mean (**Figure 3D**) and noise analysis (**Figure 3E**) reveals that ∼18% of genes are noise-regulated by SON, while only 8% appear mean-regulated. Specifically, 1,482 transcripts display higher noise in the presence of SON (i.e., control) than in the absence (i.e., KD) (**Figure 3F**), with minimal effects on their mean mRNA expression levels (**Figure 3F**, inset). These data confirm SON’s noise enhancing capabilities, and show that *Son* KD uncovers more enhanced transcripts likely because of the longer depletion time (24 hours versus 3-hour CHX treatment). Surprisingly, an additional 1,658 transcripts show lower noise in the presence of SON (i.e., control) than the absence (i.e., KD) (**Figure 3G**), indicating that SON can also repress noise of some transcripts. Since the noise repression by SON only becomes evident after longer more specific perturbations (24 hours of *Son* KD compared to 3 hours of CHX treatment), it is likely that the shorter proteome-wide CHX treatment is not sufficient to reveal noise-repressed transcripts. Alternatively, the longer 24-hour Son KD might reveal more downstream targets affected by noise propagation through key regulators that SON directly interacts with.

**Figure 3.**
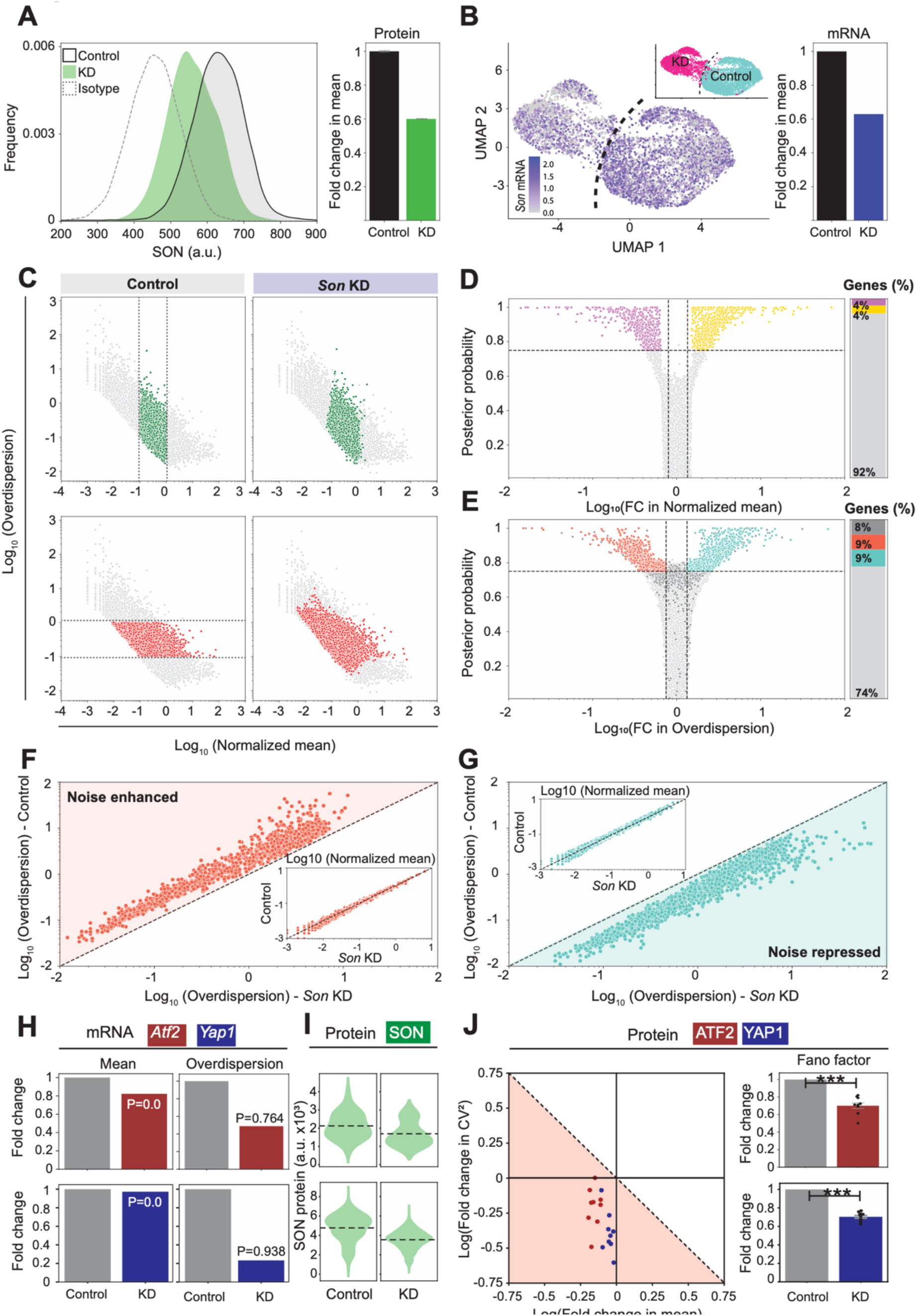
SON depletion changes noise with minimal effects on mean mRNA levels. **(A)** Left: Distribution of single-cell intensities for SON immunostaining in negative control (black) and *Son* KD (green), and isotype antibody control (grey). Right: Fold change in mean SON expression in KD compared to negative control. Error bars represent the standard error of the mean. **(B-G)** scRNA-seq of mESCs transfected for 24 hours with negative control (black) and SON siRNA (3965 and 1978 cells respectively). (B) Right: UMAP dimensionality reduction of sequenced cells after *Son* KD (inset, pink) and negative control (inset, turquoise). *Son* mRNA expression per cell is shown in blue. Left: fold change in mean *Son* mRNA compared to negative control. (C) Normalized mean expression versus overdispersion with highlighted genes displaying a central range in mean expression (top, green) or overdispersion (bottom, red) in the control population (left) and *Son* KD population (right). (D-E) Differential expression analysis with BASiCs of the *Son* KD sample compared to the control. Volcano plot represents the fold change in mean (D) or overdispersion (E) versus the certainty (Posterior probability) that this change is significant (dashed lines are the significance thresholds). Stacked bar plot represents the % genes that display significant changes in mean (D; yellow: 786 mean-repressed genes; violet: 664 mean-enhanced genes) or overdispersion (E; orange: 1482 noise-enhanced genes; turquoise: 1658 noise-repressed genes; dark grey: 1451 genes excluded because they display differential mean expression). (F-G) scRNA-seq of control population *versus Son* KD population (turquoise or orange). Overdispersion and mean expression (inset) in control *versus Son* KD populations of 1482 genes classified as noise-enhanced (F) and 1658 genes classified as noise-repressed (G). These genes change in noise levels (F-G) without changing in mean mRNA expression levels (F-G, inset). **(H-J)** Changes in *Atf2* and *Yap1* expression mean and noise upon *Son* KD. (H) Fold change in mRNA mean (left) and overdispersion (right) in *Son* KD sample, from scRNA-seq data (Figure 4), compared to negative control, for *Atf2* (top) and *Yap1* (bottom). (I) Distribution of SON expression in the control (left) and *Son* KD sample (right) for the ATF2 (top) and YAP1 immunofluorescence experiments (bottom). (J) Left: Scatter plot of Log(Fold change) in mean protein expression and CV^2^ when comparing the negative control to *Son* KD, each dot represents a randomized subset of 100 (ATF2, red) or 200 (YAP1, blue) cells. Orange highlighted area represents changes in CV^2^ that are lower than the expected changes based on Poissonian scaling of the mean. Right: Fold change in Fano factor (noise) as quantified from immunostaining in negative control compared to *Son* KD. Each dot represents randomized subset of 100 (ATF2, top) or 200 (YAP1, bottom) cells. Error bars show the standard error of the mean.

It is possible that two confounding effects influence the noise measurements upon *Son* KD: i) cell cycle changes; and ii) KD induced extrinsic cell-to-cell heterogeneity. First, an effect of SON on the cell cycle was previously described^55–57^, where *Son* KD did not stop cells from cycling, but shortened G1 and S and lengthens G2 phase. While we observed a similar trend (**Figure S4A**), the noise regulation of SON is present irrespective of the cell cycle phase of a cell (**Figure S4B-C**). Second, performing a more stringent extrinsic noise filtering step that removes heterogeneity introduced by the *Son* KD, did not have a drastic effect on the noise-regulated genes (**Figure S4D-H**). This suggests that the observed changes in noise are not dominated by variable *Son* KD between cells. While SON is not solely a noise regulator, as it also alters mean expression levels (**Figure 3D** and **Figure S4C, G)**, reanalysis of existing stochastic simulation results^18^ show that classic mean regulators––that change mean through one kinetic step––do not commonly alter noise levels without altering mean mRNA abundance (**Figure S4I**). Instead, SON solely controls noise levels of ∼18% of the transcriptome, indicating that SON is capable of noise modulation in mESCs.

In order to validate that SON’s ability to regulate gene expression noise is functionally relevant, we first confirmed that the observed effects on mRNA noise manifest at the protein level. To this end, we validated two proteins ATF2^58^ and YAP1^59–61^, which are categorized as noise-enhanced. *Atf2* mRNA shows a reduction in noise (measured as overdispersion) accompanied by a slight (non-significant) reduction in mean mRNA expression (**Figure 3H**, red) upon SON depletion. We performed immunofluorescence of ATF2 protein after *Son* KD (**Figure 3I**, top) and quantified gene expression noise after extrinsic noise filtering, which revealed a similar trend at a protein level (**Figure 3J**, red). While there is a subtle decrease in mean ATF2 levels, SON depletion decreases protein noise by 30% (**Figure 3J**, right top). Conversely, *Yap1* mRNA barely changes in mean expression levels while noise levels decrease considerably (**Figure 3H**, blue). When validating this effect at a protein level (**Figure 3I-J**), we again observed a similar behavior (**Figure 3J**, blue). Specifically, mean protein expression does not change, while noise quantified as both CV^2^ and Fano factor decrease by ∼30% (**Figure 3J**, right bottom), indicating that the noise-enhancement carried out by SON can propagate to the protein level. Notably, as this validation was not performed proteome wide it is likely that SON’s effect observed at an mRNA level does no propagate to all proteins.

### Noise-regulated genes do not change in isoform usage

To discern whether changes in splicing isoforms could account for the observed changes in noise upon SON depletion, we performed long-read RNA-seq of both 24-hour *Son* KD and control cells. Confirming the depletion of *Son* mRNA levels, a reduction of ∼3.5-fold is observable across the different transcript biotypes in which *Son* isoforms are classified (**Figure 4A**). Overall, the transcript occurrence (measured as transcript per million – TPM) in the KD correlates with the control (**Figure 4B**). Examining the distribution of biotypes across all five categories (i.e., noise/mean regulated and unchanged genes) did not reveal considerable differences following *Son* KD (**Figure 4C**). Interestingly, when focusing on isoform biotypes within the mean- or noise-regulated genes, noise-regulated genes have a higher occurrence of intron retained isoforms than both mean-regulated and unchanged genes (**Figure 4C**, green). To assess whether SON depletion influences isoform usage (**Figure S5A-D**), we calculated the absolute cumulative difference in variant usage per gene and biotype and plotted the average % difference across all transcripts when comparing the KD to the control (**Figure 4D** and **Figure S5B** and **S5D**). This reveals global changes in protein coding isoforms (both canonical and non-canonical) as well as intron retained isoforms across all five categories, likely due to the broad RNA processing functions of SON. Notably, noise-regulated transcripts do not display more striking differences compared to unchanged or mean-regulated transcripts, suggesting that observed changes in noise after *Son* KD are not associated with stark alterations in isoform usage. When considering only the dominant variant per transcript in the control sample, dominant variants that are noise-enhanced and contain a retained intron become less abundant compared to the unchanged transcripts (**Figure 4E**, orange compared to grey, and **Figure S5C-D**). Lastly, to determine if isoform usage becomes more homogenous (**Figure 4F-G**), we quantified the difference in the inverse Simpson’s index when SON is depleted. Interestingly, mean-regulated genes display more prominent changes than noise-regulated genes upon *Son* KD (**Figure 4H**). Collectively, the long-read RNA-seq data indicate that SON depletion does not alter the isoform usage of noise-regulated genes more readily than mean-regulated or unchanged genes.

**Figure 4.**
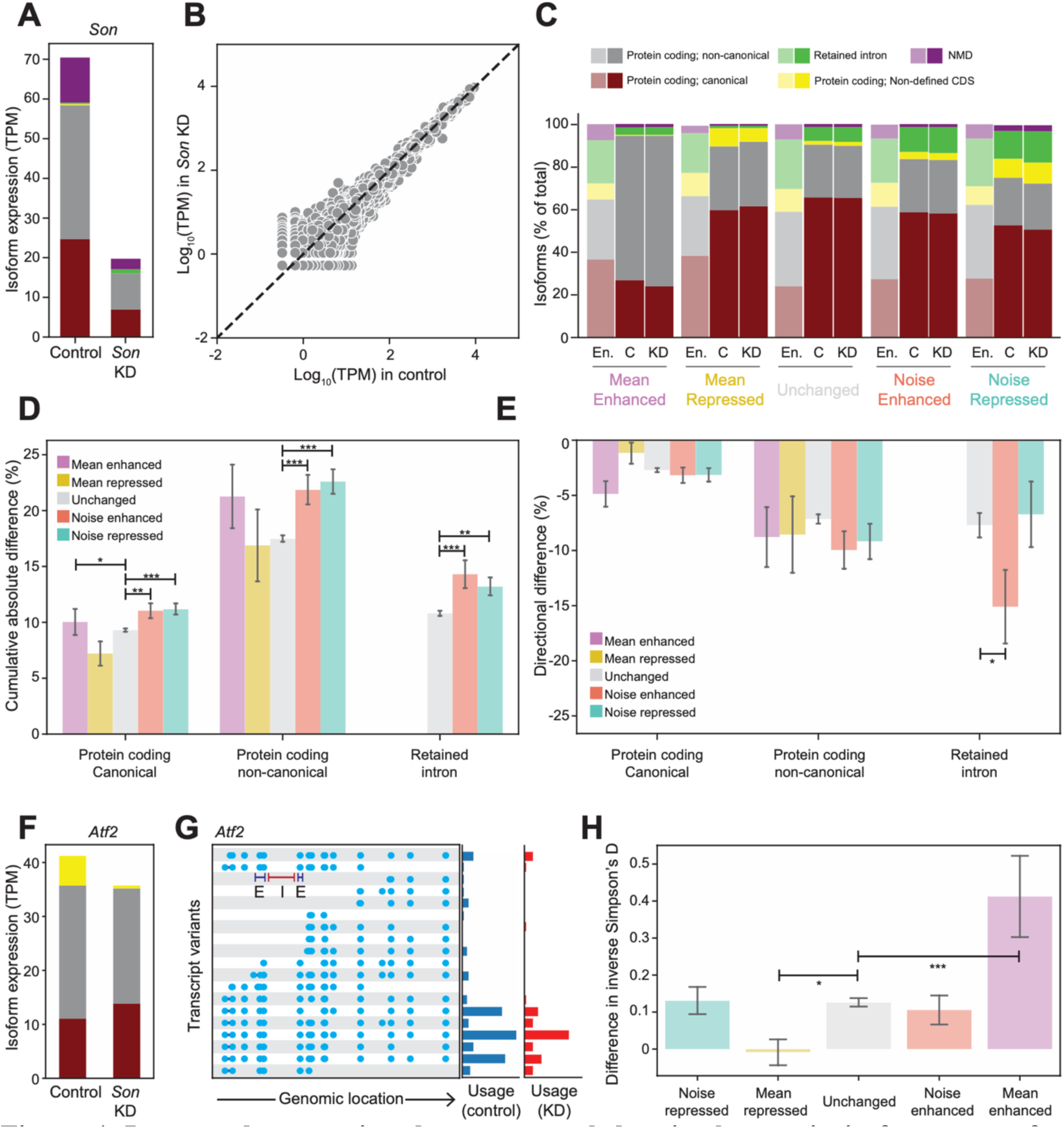
Long-read sequencing does not reveal drastic changes in isoform usage for noise-regulated genes after *Son* KD. **(A)** Quantification of *Son* transcripts from long-read RNA-seq. Read counts are classified per transcript biotype (see legend in C) and normalized to the total size of the sequenced library, denoted as transcripts per million (TPM). **(B)** Quantification of all transcript variants (TPM) in the control versus Son KD samples from long-read RNA-seq. **(C)** Distribution of transcript biotypes for all transcripts in the Ensemble database (En.) and weighted based on their read counts in the control (C) and *Son* KD (KD) samples. Genes are classified in 5 categories (mean-enhanced, mean-repressed, unchanged, noise-enhanced and noise-repressed) based on the results of the scRNA-seq analysis of control and *Son* KD populations (see Figure 4). **(D-E)** Average changes in transcript isoform usage for genes classified as mean-enhanced (violet), mean-repressed (yellow), noise-enhanced (orange), noise-repressed (turquoise) or unchanged (grey) upon *Son* KD, as defined by the scRNA-seq analysis. (D) Cumulative absolute differences in usage for all variants (|% in KD - % in control|). (E) Directional differences in usage quantified for the most common variant of each gene in the control situation (% in KD - % in control). Transcript variants are classified according to their biotypes. Biotypes with a representation of <5% in the unchanged population or the respective mean/noise groups were discarded from the analysis. (F) Quantification of *Atf2* transcripts from long-read RNA-seq. Read counts are classified per transcript biotype and normalized to the total size of the sequenced library, denoted as TPM. (G) Representation of *Atf2* transcript isoforms in Ensembl database (left). Blue lines represent exons, while gaps between exons represent introns. Exons and introns are located based on their respective genomic locations. Bars (right) represent variant usage (% of total gene TPM) in control sample (blue) and *Son* KD (red). (H) Change in isoform diversity calculated by the difference in Inverse Simpson’s D in the *Son* KD sample compared to the control for genes classified as mean-enhanced (violet), mean-repressed (yellow), noise-enhanced (orange), noise-repressed (turquoise) or unchanged (grey). (D, E, H) Error bars represent standard error of the mean change in inverse Simpson’s D. Significance is computed with two-tailed independent samples t-test and indicated with * (p-value <0.05), ** (p-value<0.005) and *** (p-value<0.0005).

### Noise regulation is not associated with discernable differences in splicing efficiency

Considering that SON noise regulation does not appear to act by altering isoform usage, we instead examined whether SON alters noise by modulating splicing efficiencies. We thus performed total RNA-seq after rRNA depletion and quantified splicing efficiency of introns^62^ on *Son* KD and control cells. The data reveals an enrichment in introns that decrease in splicing efficiency (SE FC < 10%) in the group of mean enhanced genes upon *Son* KD (**Figure 5A**, violet), consistent with SON’s role in enhancing weak splicing sites.^63^ Furthermore, the group of mean repressed genes also show an enrichment in introns whose splicing efficiency increased when SON is depleted (**Figure 5A**, yellow), suggesting that SON is capable of reducing splicing efficiency of some transcripts in mESCs. Interestingly, noise-regulated genes display comparable splicing efficiency to the background group of mean and noise unchanged genes (**Figure 5A**, orange and turquoise). It therefore appears that SON alters mean mRNA levels through changing splicing efficiency, while noise modulation either does not occur through changes in splicing efficiency or changes in splicing efficiency are too subtle to detect.

**Figure 5.**
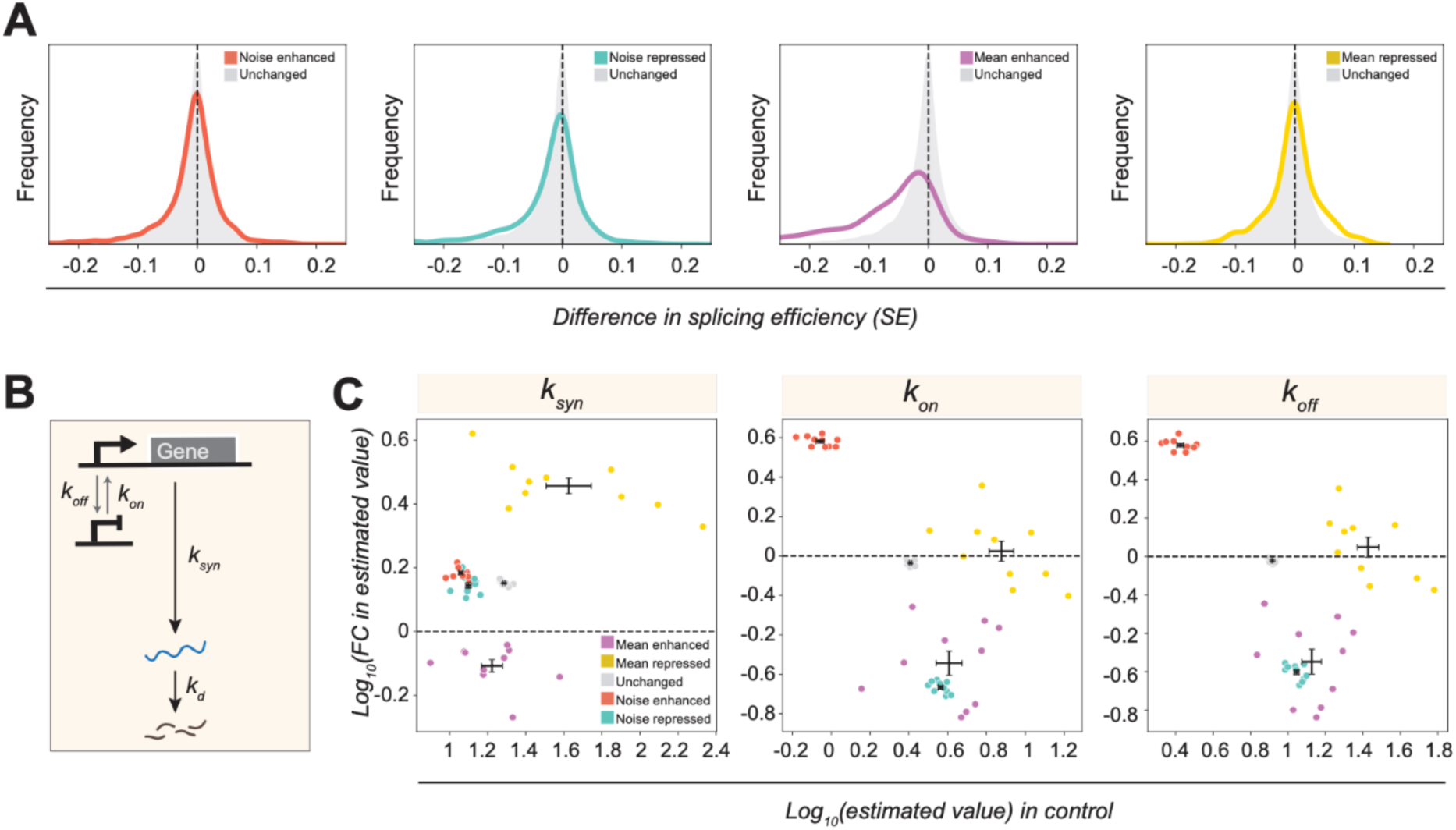
Noise regulation by SON is not linked to splicing efficiency. **(A)** Distribution of changes in splicing efficiency (SE) upon *Son* KD, for single introns from genes classified as (from left to right) noise enhanced (orange), noise repressed (turquoise), mean enhanced (violet) and mean repressed (yellow). Grey distributions in every plot represent genes classified as unchanged in mean and noise. Distributions correspond to the average of two replicates. **(B)** Schematic of the two-state random-telegraph model. **(C)** Estimation of parameters of the model in B from the scRNA-seq data in Figure 3 in +SON conditions (control sample) and fold change in *Son* KD versus the control sample. From left to right: *k_syn_*, *k_on_*, and *k_off_*. Each point represents a subsampled set of 10% of genes corresponding to each mean/noise modulation category.

Next, we hypothesized that if SON’s involvement in splicing is co-transcriptional, then it might impact transcription or at least the apparent transcription rates obtained from the scRNA-seq data.^64,65^ We therefore utilized a neural-network based approach to fit the scRNA-seq data (**Figure 3**) to a two-state random telegraph model (**Figure 5B**) to extract effective rates for *k_on,_ k_off_, k_syn_*.^66^ We computed the average rates in the control as well as the average change in rate after *Son* KD for 10 sub-sampled groups consisting of 10% of all genes in each category (i.e., noise/mean regulated and unchanged genes). Notably, the estimation of these rates is with respect to mRNA degradation, which are assumed not to be affected by SON depletion. *Son* KD does not alter *k_syn_* of noise regulated genes (**Figure 5C**, left) compared to unchanged genes. Conversely, SON depletion increases *k_syn_* of mean repressed genes and decreases *k_syn_* of mean enhanced genes (**Figure 5C**, left), which is consistent with the splicing efficiency analysis. Given that the estimated *k_syn_* will likely be dominated by either transcription or splicing–– depending on which of the two steps is rate limiting––it is intuitive that SON can alter mean gene expression levels through *k_syn_*.

On the other hand, SON depletion causes both *k_on_* and *k_off_* (**Figure 5C**, middle and right) to increase for noise enhanced genes and decrease for noise repressed genes compared to noise unchanged genes. Notably, the underlying model is an oversimplification and does not capture the mechanism by which SON acts. Therefore, this analysis only indicates that SON changes mean mRNA levels by altering splicing efficiency, while noise modulation likely acts through an alternate mechanism. For example, it is possible that: i) SON’s role in splicing or nuclear speckle association impacts transcriptional bursting ^67,68^; or ii) that mRNA processing by SON is in itself a bursty process.

### Tubercidin treatment alters mRNA noise revealing commonalities and distinctions from SON depletion

We next considered that SON’s noise regulation might be connected to its role in nuclear speckle formation.^25,68^ We therefore treated mESCs with a small-molecule disrupter of nuclear speckles––Tubercidin^69^––which SON has been shown to associate with.^24–26^ Tubercidin, is an adenosine analogue that has been shown to disrupt nuclear speckles. Since it is a base analogue, it could affect RNA metabolism independently of speckle disruption. Therefore, to limit downstream effects of Tubercidin treatment, we only treated for short time periods (3 hours for RNA and 6 hours for protein quantification.

We first treated mESCs with 10 µM of Tubercidin for 6 hours and imaged speckles through SON immunofluorescence in both the DMSO control and Tubercidin treated cells (**Figure 6A**). SON signal is classified as a speckle if it is brighter than a threshold set according to randomized signal (see STAR Methods for more information) and between 0.23-2.33 µm^2^. We subsequently quantified average SON intensity per cell, number of speckles per cell, as well as average speckle size and intensity per cell (**Figure 6B**). Tubercidin treatment reduces both SON intensity quantified over the entire nucleus and speckle abundance per cell (**Figure 6B**, top), while speckle size and intensity remain more similar (**Figure 6B**, bottom). The latter can be explained either by the analysis method, since we define everything above a certain size and SON intensity as a speckle, or because in order to form a speckle a critical amount of SON is required. Therefore, short Tubercidin treatment might disrupt some speckles––reflected in 40% fewer speckles and a decrease in nuclear SON intensity––while remaining speckles contain similar SON levels as the control––reflected in their size and intensity. Finally, as we classify speckles solely through SON staining, while all speckles are expected to contain SON it is possible that not all quantified speckles are true nuclear speckles.

**Figure 6.**
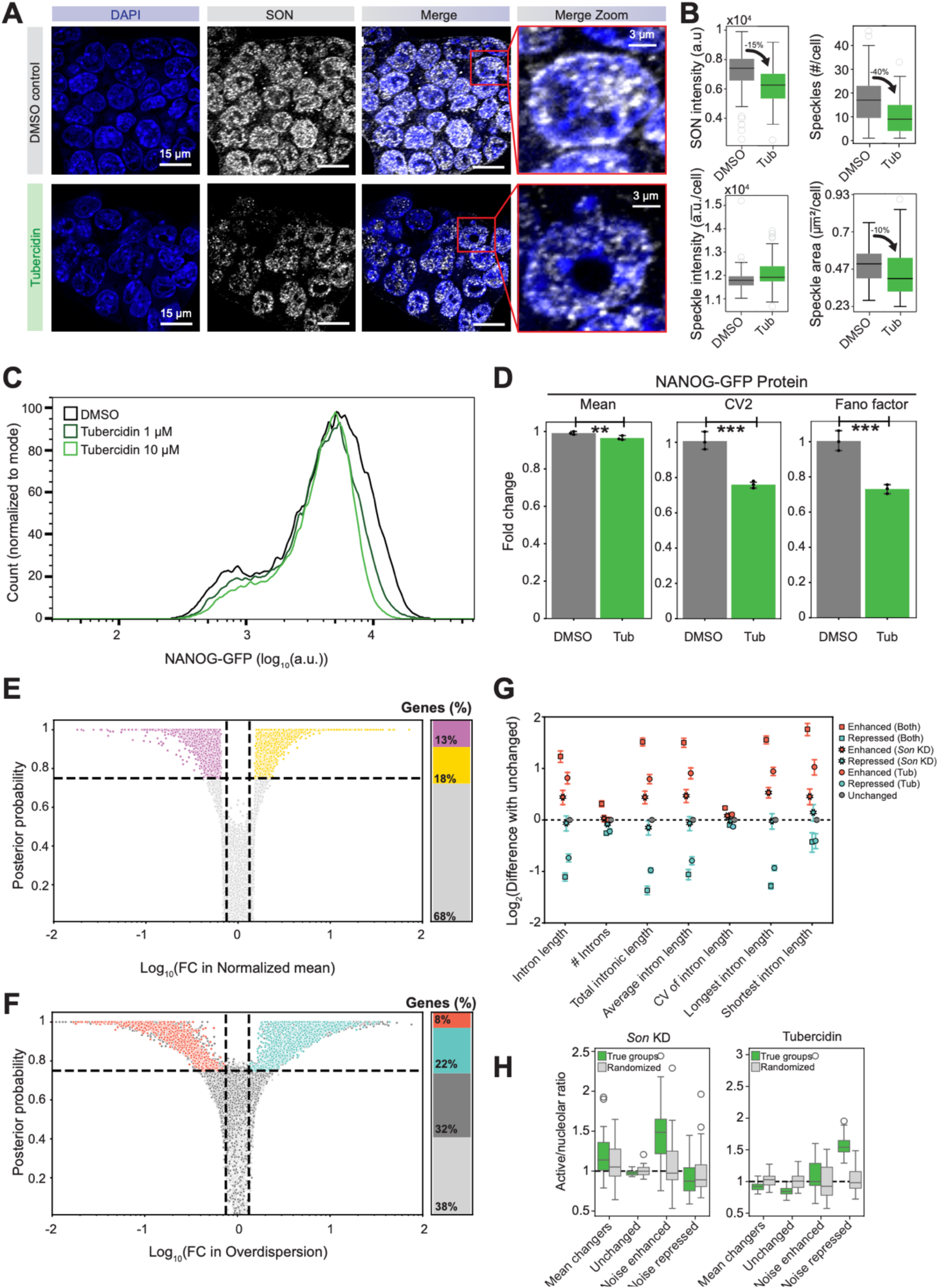
Tubercidin-driven disruption of nuclear speckles reveals commonalities and distinctions from SON depletion. **(A-B)** Disruption of nuclear speckles with Tubercidin treatment. (A) Representative microscopy images of mESCs from DMSO control (top) and Tubercidin treatment (bottom) in the DAPI channel, SON immunostaining, merged, and zoom in of one cell (left to right). (B) Quantification of single-cell SON intensity, number of detected speckles per cell, average intensity of detected speckles per cell and average area of detected speckles per cell over 2085 speckles detected in 123 cells in control conditions and 1109 speckles detected in 108 cells in *Son* KD conditions. **(C-D)** Flow cytometry analysis of NANOG-GFP mESCs. (C) Distribution of NANOG-GFP intensity for DMSO control (black), 1 µM (dark green), and 10 µM (light green) Tubercidin treatment for 6 hours. (D) Fold change in mean NANOG-GFP protein (left), CV^2^ (center), and Fano factor (right) upon treatment with 10µM Tubercidin for 6 hours compared to control after extrinsic noise filtering. Each dot represents a biological replicate. Error bars show 95% confidence intervals. **(E-F)** Differential expression analysis with BASiCs of scRNA-seq data from mESCs treated with 10 µM Tubercidin for 3 hours compared to the control (1968 and 2594 cells respectively). Volcano plot represents the fold change in mean (E) or overdispersion (F) versus the certainty (Posterior probability) that this change is significant (dashed lines are the significance thresholds). Stacked bar plots represent the % genes that display significant changes in mean or overdispersion (violet: 2264 mean-enhanced genes; yellow: 3096 mean-repressed genes; orange: 1302 noise-enhanced genes; turquoise: 3813 noise-repressed genes; dark grey: 5360 genes excluded because they display differential mean expression). **(G)** Intron characteristics of genes that are noise enhanced (orange) or noise reduced (turquoise) in the *Son* KD (asterisks), Tubercidin treatment (circles), or both (squares). Marker represents the average value for 10 randomized samples of 100 genes in a given group and for the indicated characteristic, normalized to the group of unregulated genes. Error bars represent the standard error of the means. **(H)** Physical association with nuclear speckles of genes from different noise/mean regulation categories from either *Son* KD (left) or Tubercidin (right). The association with nuclear speckles is defined by the active/nucleolar ratio as previously obtained,^74^ in 20 random groups of 50% of genes from each category. Grey boxes correspond to randomized groups of genes of the same size. Dashed line represents a similar association with active and nucleolar compartments.

We next hypothesized that if SON’s noise regulation mechanism is associated with nuclear-speckle formation, disrupting these speckles should also impact noise without altering mean protein expression levels. Since *Nanog* shows no significant change in mean mRNA or noise when SON is depleted, we expected that we would only see an effect of Tubercidin on *Nanog* expression if chemically disrupting nuclear speckles is more potent or acts on a different subset of transcripts than knocking down SON. Strikingly, Tubercidin considerably reduces NANOG noise (quantified in a NANOG-GFP expressing mESC line^70^) in a dose-dependent manner (**Figure 6C-D**) with minimal effects on mean expression levels (**Figure 6C-D**) and cell viability (**Figure S6A** bottom, inset). Importantly, this noise reduction occurs even in the presence of extrinsic noise (**Figure S6B**), indicating that the noise modulation resulting from nuclear speckle formation is not overridden or reversed by extrinsic noise sources. These data suggest that SON’s noise regulation could act––at least in part––through the association with nuclear speckles.

To further explore the commonalities and distinctions between nuclear speckle disruption and SON depletion, we produced scRNA-seq data from Tubercidin treated cells (10 µM, 3 hours) and measured the transcriptome-wide changes in mean and noise (**Figure 6E-F**). Tubercidin treatment has a stronger impact on mean expression compared to SON KD (**Figure 6E** compared to **Figure 3D**). Furthermore, the data show that nuclear speckle disruption through Tubercidin reveals more noise suppression (22%, **Figure 6F** turquoise) than SON depletion, while noise enhancement is still occurring (8%, **Figure 6F** orange). This is consistent with previous reports where membrane-less compartments have been shown to suppress noise.^71^

Since Tubercidin treatment likely causes other effects not related to nuclear speckle disruption it does not entirely mimic SON depletion. We therefore sought to determine whether certain transcript properties might suggest shared noise regulation aspects. Thus, with the single-cell data from both *Son* KD and Tubercidin treatment, we attributed each gene to a shared category in terms of noise regulation and analyzed transcript properties of these categories. We next quantified the intron length––which is thought to be associated with mRNA processing efficiencies^72,73^––of genes within these different categories. Interestingly, noise enhancement is associated with longer introns and intronic length in *Son* KD, Tubercidin treatment, and in the shared enhanced categories (**Figure 6G**, orange asterisks, circles and squares). This indicates a potential common mechanism of noise enhancement––associated with nuclear speckles––that might be impacted by both reduced SON levels and by Tubercidin treatment. Conversely, introns of noise repressed genes in Tubercidin treatment and in shared genes (i.e., both Tubercidin and *Son* KD) are shorter in length (**Figure 6G**, turquoise circles and squares). While noise repressed genes in the *Son* KD data behave similar to the unchanged group (**Figure 6G**, turquoise asterisks compared to grey circles), genes that are noise repressed in both the *Son* KD and Tubercidin treatment display shorter intron length than genes that are solely noise repressed in the Tubercidin data. This implies that either two distinct mechanisms explain the noise repression observed in *Son* KD and Tubercidin, or that differences in intron length are too subtle to distinguish in the *Son* KD only category. To explore which is more likely, we plotted the distribution of intron lengths for noise enhanced, repressed and unchanged genes. These data reveal that while the shift in the distribution of noise enhanced genes towards longer introns is evident in the *Son* KD only category (KS test p =1.8×10^-56^; **Figure S6D**), there is also a subtle but significant (KS test p = 1×10^-13^) change in the distribution of noise repressed genes in the *Son* KD only category (**Figure S6E**). On the other hand, the exon length distributions remain unchanged (**Figure S6D-E**, insets).

Finally, we sought to explore the link between noise regulation and physical association of genes with nuclear speckles. We thus exploited previously reported classifications,^74^ where genes were classified as speckle-associated (i.e., active compartment) or speckle-anti-associated (i.e., nucleolar compartment). Intriguingly, noise enhanced genes from the *Son* KD and noise repressed genes from the Tubercidin treatment display increased association with the active compartment––i.e., nuclear speckles (**Figure 6H**). Notably, it is possible that the limited overlap between Tubercidin treatment and Son KD is because Tubercidin can alter other RNA processing steps that SON is not involved in, or because of differences in perturbation times (24 and 3 hours respectively). Together, these data indicate that while perhaps not all noise regulation by SON can be explained by nuclear speckle association, nuclear speckles could play a role in noise regulation in cells.

### SON depletion impacts cell-fate choices

Finally, we sought to determine SON’s involvement in differentiation, a process known to be affected by gene expression noise.^11^ In particular, high noise is beneficial as cells exit pluripotency, to allow for cells to differentiate down separate lineages.^8,9^ Gene ontology enrichment analysis revealed that noise regulated genes are related to differentiation, development or morphogenesis, with a higher percentage represented in noise enhanced genes (**Figure 7A, Table S2**). Therefore, we determined the effect of SON depletion on differentiation of mESCs, utilizing CD24 as a marker for differentiation.^11,75^ We observed a weak but significant decrease in differentiation efficiency (i.e., % CD24+ cells) quantified 96 hours into differentiation (**Figure 7B** and **Figure S7A-C**). These results differ from previous experiments in human ESCs (hESCS), where SON depletion results in increased expression of differentiation markers in undifferentiated cells.^27^ These differences could be due to the cell-type, culture, and markers used, or the longer depletion time of SON and correspondingly higher cell death in previous studies.^27^ Furthermore, the differentiation potential of hESCs upon SON depletion was not assessed in this previous study. Based on our data, we hypothesized that SON’s noise enhancement was likely more relevant during exit from pluripotency than noise repression. Moreover, if SON is involved in noise enhancement, then introducing an excess of noise during SON depletion should push the system into the opposite direction. To test this, we treated SON depleted cells with a published small-molecule noise enhancer (5-Iodo-2’-deoxyuridine, IdU),^11^ and measured differentiation efficiency. Markedly, in the absence of SON and in the presence of the noise enhanced control differentiation is significantly increased.

**Figure 7.**
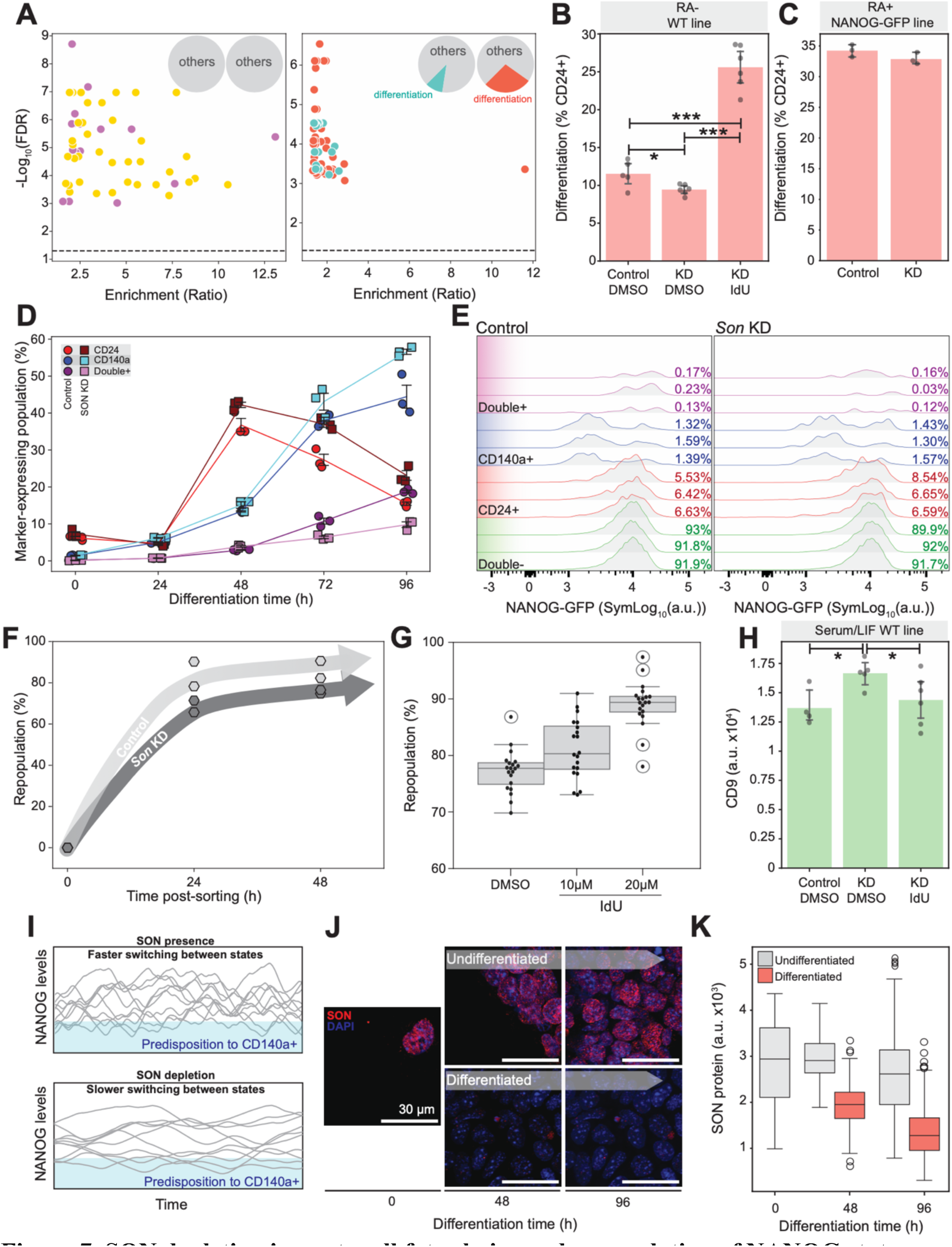
SON depletion impacts cell fate choice and repopulation of NANOG states. **(A)** Gene Ontology enrichment analysis for genes identified as mean (left) enhanced (yellow) or repressed (violet), or noise (right) enhanced (orange) or repressed (turquoise), from the analysis of *Son* KD scRNA-seq data. Pie charts (insets) represent the ratio of significantly enriched Gene Ontology terms that are related to differentiation, development or morphogenesis, *versus* other terms. **(B-C)** Differentiation efficiency quantified as percent of cells expressing the membrane differentiation marker CD24 after 96 hours of differentiation, in wild type mESC in control, *Son* KD and *Son* KD + 4 µM IdU samples and in absence of retinoic acid. Error bars show 95% confidence intervals of 6 replicates. **(B-C)** Differentiation efficiency quantified as percent of cells expressing the membrane differentiation marker CD24, in NANOG-GFP mESCs and in presence of retinoic acid. Error bars show 95% confidence intervals of 3 replicates. **(D)** Fate choice during differentiation of NANOG-GFP mESCs in control (circles) and *Son* KD (squares), quantified as percent of cells expressing the membrane differentiation marker CD24 (red/maroon), the membrane differentiation marker CD140a (blue/cyan), or both (purple/violet). Measurements were performed every 24 hours during the differentiation process, with 3 replicates per condition and timepoint. Error bars represent the standard error of the mean of each replicate. **(E)** Distributions of NANOG-GFP signal at 0 hours of differentiation, for control cells (left) and 24-hour *Son* KD (right), for cells expressing the CD24 marker (red), the CD140a marker (blue), both (purple), or none (green). **(F)** Repopulation (%) of NANOG-GFP mESCs sorted, into NANOG-high and NANOG-low populations, and grown for 24 and 48 hours post-sorting, in control (light grey) and *Son* KD (dark grey) conditions. Each hexagon marker represents the mean of 20 randomized subsets of 1000 cells for each biological replicate. **(G)** Repopulation (%) in the presence of 10-20 µM of IdU between from 24 hours to 48 hours the *Son* KD condition. Each point represents the mean repopulation for a randomized subset of 1000 cells. **(H)** Immunostaining and flow cytometry analysis of the expression of membrane pluripotency marker CD9 in mESCs in DMSO control, *Son* KD or *Son* KD + IdU samples. Error bars show 95% confidence intervals of 6 replicates. **(I)** Schematic of cells demonstrating fast switching (top; SON presence) and slow switching (bottom; SON depletion) between NANOG states. Blue shaded area represents predisposition to CD140a fate when cells are expressing low NANOG levels. **(J-K)** SON expression during differentiation of mESCs. (J) Representative confocal microscopy images of SON at different time points of differentiation. Blue represents nuclear DAPI staining and red represents SON immunofluorescence. (K) Single-cell quantification of SON expression during differentiation (red) or undifferentiated control (grey). In B and H, significance is computed with one-tailed independent samples t-test and indicated with *(p-value<0.05), ** (p-value<0.005), and *** (p-value<0.0005).

Given the weak impact of SON depletion on differentiation efficiency, we considered a different aspect of differentiation not linked to the efficiency of the process, but to the stability of the cell fate choice. To this end, we explored how SON depletion impacts the choice between the neuroectodermal fate (CD24+) and the extraembryonic endoderm fate (CD140a+).^75^ To link the ultimate choice to cellular predisposition, NANOG-GFP mESCs were employed. The stability of the two different fates was measured as % of cells presenting exclusively CD24 or CD140a markers, while Double+ cells were interpreted as unstable fate exploration. Again, *Son* KD did not drastically alter the global CD24+ differentiation (**Figure 7C**), yet stable differentiation in both ectoderm and extraembryonic endoderm fates increased and Double+ (i.e., unstable) cells were reduced, at 96h of differentiation (**Figure 7D** and **Figure S7D**). To determine if SON depletion impacted cell-fate predisposition, NANOG-GFP levels were quantified in CD24+ and CD140a+ cells at 0 hours. Interestingly, some NANOG-low cells showed increased CD140a+, indicating early signs of predisposition to the extraembryonic endoderm fate (**Figure 7E** and **Figure S7D**). Importantly, SON depletion does not appear to impact the percentage of cells predisposed to a particular fate (**Figure 7E**, see percentages).

We therefore hypothesized that SON’s noise enhancement could be linked to exploration of fates rather than altering a cell’s predisposition to a fate. To validate this hypothesis NANOG-GFP cells were sorted into NANOG-high and NANOG-low populations and the repopulation was measured for *Son* KD and control cells. Notably, this repopulation will depend on cells’ ability to switch from the NANOG-low to the NANOG-high state or *vice versa*, where faster repopulation indicates faster switching. After SON depletion, repopulation is slowed down, indicating that the presence of SON allows for faster NANOG-state switching (**Figure 7F**). Furthermore, treatment of SON depleted cells with the noise enhanced (IdU) control increases the repopulation in a concentration dependent manner (**Figure 7G**), showing that the reduced NANOG-state switching when SON is removed is enhanced in the presence of increased noise. These results were confirmed in the mESC line by using CD9 as a marker for pluripotency.^76^ Specifically, 48 hours of SON depletion significantly increases CD9 levels, while the noise enhanced (IdU) control brings the levels of CD9 back down to the control sample (**Figure 7H** and **Figure S7E-G**). These data indicate that the presence of SON allows for faster switching between NANOG-low and NANOG-high states and thus favors an exploration of different fates **(Figure 7I**). Finally, throughout 96 hours of differentiation a gradual decrease of SON was observed (**Figure 7J-K**), implying that SON’s role is more important during (exit from) pluripotency than during differentiation.

## DISCUSSION

Here we perform a multimodal screen in mESCs and discover 32 endogenous proteins that could be involved in noise regulation. In contrast to some previous studies,^12,77^ our strategy uses direct single-cell transcriptomic measures of noise. Furthermore, our approach is not based on the perturbation of one specific molecular process to study changes in noise,^12,18,22,39,78^ but rather targets the full proteome and reveals a novel class of proteins as noise-regulators in a mechanism-unbiased manner. This study makes use of pre-existing knowledge by exploiting previously described regulator-target associations in the regulator enrichment analysis. Thus, as more data become available connecting regulatory proteins to mRNA targets, the list of noise regulators will likely expand using the methodology developed and data reported herein. However, target-regulator relationships that are classically defined based on mean gene expression levels (e.g., mRNA correlation^42^) were used to identify noise regulation. Therefore, more noise regulators will likely emerge if in future the regulator-target relationship is also defined by changes in noise.

The reported findings indicate that proteins might have evolved to regulate noise orthogonally to mean mRNA and protein expression levels, distinguishing them from classic mean repressors and activators. The discovered proteins could play an important role in cellular noise-control to ultimately impact cell-fate transitions, which is relevant for our understanding of developmental processes^79^ as well as pathogenesis.^3,28^ We find that the noise regulator SON, alters noise with minimal effects on mean mRNA expression levels of ∼3,100 genes. Since scRNA-seq is performed after 24 hours of Son KD, we cannot distinguish between targets with which SON directly interacts with or targets that change in noise due to noise propagation. Currently, existing databases define regulator-target relationships relatively loosely. Therefore, if new data emerges that defines these relationships more stringently, it will likely become possible to make these distinctions based on the gene-lists provided herein.

While it appears that SON’s ability to regulate noise is mechanistically distinct from mean regulation, the exact mechanism through which noise is regulated remains to be elucidated. Given that SON is a pleiotropic regulator, uncovering the exact mechanism through which SON alters noise directly (i.e., through its direct targets) or indirectly (i.e., through noise propagation), might be a complex process. The dual role of SON in regulating both noise and mean expression suggests that regulatory proteins may have evolved to control noise as well as the mean expression levels of their targets. Yet, this does not necessarily imply that noise regulation is more important than mean expression regulation by a protein.

The implications of SON as a noise-regulator could exceed the limits of the field of stem cells, as this protein has been associated with neurological and syndromic disorders.^80^ Furthermore, liquid-like compartments have been linked to noise reduction,^71^ and our finding that nuclear speckle formation alters gene expression noise suggests a broader role of membrane-less organelles in noise control. The developed screen––as well as the discovered small-molecule noise-modulator, Tubercidin––could provide a valuable resource for future studies aimed at linking noise control to cell-state transitions^81^ in various biological systems.

Previous research has shown that across the kingdoms of life noise is harnessed or suppressed by cells depending on the biological context. Classic examples include pathogenic systems that harness noise to facilitate state-transition––from a drug susceptible to tolerant state––to enhance fitness. These examples span bacterial antibiotic persistence, viral infections, or drug tolerance in cancer cells.^5,6,82,83^ On the other hand, a striking example where noise repression is enforced is developmental patterning in the early *Drosophila* embryo.^14^ Overall, cell fate commitment requires low noise because cell-state transitioning is detrimental^84^ and cell fate diversification requires high noise as cell-state transitioning is necessary.^82^ This highlights the importance for cells to be able to accurately modulate noise and implies that noise-control strategies might be ubiquitous. Noise-control through protein regulators would be an efficient strategy to up- or down-regulate noise simply by controlling regulator abundance. Thus, evolution could have shaped mechanisms through which genes or gene-subsets could be temporally switched from high to low noise (or *vice versa*) depending on the context.

Notably, the cell type used herein or in any future screen would likely strongly influence the outcome. For example, in systems where noise facilitates state transitions and high noise levels are advantageous it is likely that more noise enhancing mechanisms are uncovered. Conversely, during cell fate commitment or precise developmental processes where noise is unfavorable noise reducing mechanisms are likely more prominent. Consequently, the prevalence and type of noise regulators identified is expected to vary significantly depending on the specific cellular context used in the screen.

In summary, this is the first study to identify a class of proteins that regulate noise without changing mean expression levels of target mRNAs. We describe a methodology that will allow identification of regulators of gene expression noise in various physiological (i.e., development) and pathological (i.e., cancer progression) systems. In future, this could lead to new therapeutic approaches for diseases where gene expression noise plays a critical role._77,83,85,86_

## Acknowledgments

We thank Tom de Greef, Hans Heus, and members of the Hansen lab for thoughtful discussions. We are very grateful to Leor Weinberger for valuable feedback on the manuscript, as well as Xinyue Chen and Binyamin Zuckerman for their suggestions. We also thank Hendrik Marks for the mESC-E14 cell line; Leor Weinberger for the NANOG-GFP mESC line; Laura Wingens, Teun Pingen and Klaas Mulder from the RIMLS Sequencing Facility for technical assistance with scRNA-seq. M.M.K.H acknowledges generous support from the Christine Mohrmann fellowship; the Netherlands Organization for Scientific Research (NWO) [OCENW.XS3.055 and VI.Vidi.223.065]; and the European Union (ERC Starting Grant, ChOICE, 101041939).

## Author contributions

Conceptualization: OGB, MMKH

Methodology: OGB, MMKH

Investigation: OGB, XH, CLW, TvB, FMBS

Visualization: OGB, MMKH

Funding acquisition: MMKH

Project administration: OGB, MMKH

Supervision: MMKH

Writing – original draft: OGB, MMKH

Writing – review & editing: OGB, XH, MMKH

## STAR METHODS

### RESOURCE AVAILABILITY

#### Lead contact

Further information and requests for resources and reagents should be directed to and will be fulfilled by the lead contact, Maike M.K. Hansen.

#### Data and code availability

Single-cell, bulk and long-read RNA sequencing data have been deposited at Gene Expression Omnibus (GEO) and are publicly available as of the date of publication (GSE272056, GSE271660, GSE270846, GSE271659, GSE271082). Accession numbers are also listed in the key resources table. Proteomics data from LC-MS/MS experiment is provided as supplementary datasets.

Any additional information required to reanalyze the data reported in this paper is available from the lead contact upon request

### EXPERIMENTAL MODEL AND STUDY PARTICIPANT DETAILS

#### Cell lines

The majority of experiments in this publication have been performed utilizing E14 mESCs (129/Ola background), obtained from Hendrik Marks, originally from ATCC^87^, with RRID:CVCL_9108. Selected experiments were performed with NANOG-GFP E14 mESCs, provided by Leor S. Weinberger ^11^. Both cell lines are male cell lines. In both cases, mESCs were routinely cultured in gelatin (Sigma-Aldrich Cat#48723/500g) coated 9.6 cm2 wells in 6-well Clear TC-treated Multiple Well Plates (Costar® REF3516). mESC culture medium (serum/LIF) consisted of Dulbecco’s Modified Eagle Medium (DMEM; GibcoTM Cat#11995065), supplemented with 0.1mM beta-mercaptoethanol (Fisher Scientific Cat#11528926), 1000 U/mL of Penicillin, 0.1 mg/mL of Streptomycin (Fisher Scientific Cat#15070063), 1mM of sodium pyruvate (Fisher Scientific Cat#11530396), 1000 U/mL of ESGRO recombinant Mouse Leukemia Inhibitory Factor (LIF; Merckmillipore ESG1107) and 15% of ES-qualified heat inactivated Fetal Bovine Serum (FBS; Fisher Scientific Cat#A3840002). Cells were grown in a 37 °C incubator with 5% CO2 atmosphere. Cells were routinely passaged after two days of growth following a 1:6 dilution.

### METHOD DETAILS

#### Translation inhibition experiments

Cycloheximide (CHX; Sigma-Aldrich Cat#C7698) diluted in 100% Ethanol was used in order to block protein synthesis in mESCs. The stock solution of cycloheximide in 100% EtOH was 10 mg/mL. The addition of CHX in the mESC culture medium was done by mixing 7 µL of 10 mg/mL CHX solution in 100% EtOH per 1 mL of culture medium, to obtain a 250 µM working concentration. For lower working concentrations, the stock solution was previously diluted in 100% EtOH before mixing with the culture medium, so the proportion of EtOH is kept constant. Translation inhibition was achieved by changing the medium in which cells were growing 24 hours after seeding with medium containing the desired concentration of CHX. Control treatment was performed by exposing the cells to culture medium with equivalent volume of 100% EtOH but lacking CHX.

#### HPG incorporation assay

5×10^5^ mESCs were attached per well of a Lab-Tek II Chambered Coverglass W/Cover #1.5 Borosilicate Sterile Well plate (REF 155409). Cell attachment was achieved with the following procedure: First, wells were washed with water and let dry. A mixture of 20µL of Cell-Tak^TM^ (Cornig® Cat#CLs354240), 5µL of NaOH 1M and 570µL NaHCO_3_ 7.5% was prepared and used to cover the surface of the four wells (150 µL per well). After a 10-minute incubation the liquid was removed and the wells washed with water. 2×10^6^ cells were washed in 5 mL of DPBS, pelleted and resuspended in 600µL of DPBS. 150 µL of cell suspension was deposited into each well and cells were left to attach for 30 minutes. Following the attachment, the rest of cell suspension was removed and 150 µL of 250 µM CHX pre-treatment medium was added to one well. After 2.5 hours of pretreatment, medium was changed to 250 µM CHX or EtOH treatment medium in all wells. Treatment medium consisted of an L-methionine free version of mESC culture medium (substituting DMEM Cat#11995065 by DMEM Cat#21013024) and supplementing with 146 mg/L L-glutamine and 313 mg/L L-cystine, supplemented with L-homopropargylglycine (HPG; From Click-iT™ HPG Alexa Fluor™ 488 Protein Synthesis Assay Kit; Invitrogen™ Cat#C10428) except in the HPG-control. Treatment was followed by fixation with 4% paraformaldehyde in PBS for 10 minutes at room temperature, followed by two washes with DPBS and permeabilization with ice cold 70% EtOH for 1 hour at -20°C. Alexa-488 HPG detection through Click-it chemistry was performed according to the manufacturer’s instructions (Click-iT™ HPG Alexa Fluor™ 488 Protein Synthesis Assay Kit; Invitrogen™ Cat#C10428). Nuclear staining was performed as described by the manufacturer of the kit. Imaging was performed with a CSU spinning disc confocal microscope with a 40x 0.95NA objective, for alexa-488 and nuclear staining signals. Imaging of a central z-slice per position was performed with a 488 nm laser at 50% power for 300 ms and a 405 nm laser at 50% power for 100 ms, with the gain set to 200. Cells were segmented with ImageJ, utilizing the DAPI channel as input. An intensity threshold was applied to input images, to convert them to binary images. Single-cells were segmented from a binary image utilizing the watershed, fill holes and analyze particles functions from ImageJ. Produced masks where used to obtain single-cell measurements of the signal in the HGP-alexa488 channel.

#### Flow cytometry analysis of endogenous fluorescently-labelled proteins upon translation inhibition

5×10^5^ NANOG-GFP mESCs were seeded per gelatin coated 9.6 cm^2^ well (Costar® REF3516 plate). 24 hours post-treatment, cells were exposed to mESC culture media with specific concentrations of CHX of 0.2, 0.875, 2.5, 25 or 250 µM, or equivalent volume of EtOH. Cells were treated for 3,6, 9 or 12 hours, after which flow cytometry analysis of GFP signal was performed in a BD FACSCalibur Flow cytometer. Data analysis was performed with BD FlowJo.

#### Cell viability assay

Cell viability was assessed by mixing 18 µL of cell suspension with 2 µL of Acridine Orange / Propidium Iodide Stain (Logos Biosystems Cat#F23001). 10 µL of the mix were pipetted into each chamber of a LUNA^TM^ Cell Counting Slide (Logos Biosystems Cat#12001), for two technical replicates. Cell viability in each chamber was measured in a LUNA-FL^TM^ Dual Fluorescence Cell Counter (Logos Biosystems Cat#L20001). Measurements were performed in duplicates.

#### 5EU#incorporation assay

5×10^4^ mESCs where seeded per well in gelatinized µ-Slide Angiogenesis chip wells (ibidi Cat#81506). 24 hours post-seeding, cells were exposed to mESC culture medium containing CHX 250µM for 0 to 6 hours of pre-treatment. Controls exposed to corresponding volume of EtOH for 6 hours were included. Following this incubation, cells were exposed to mESC culture medium containing 0.5 mM 5EU for 30 minutes. The control for inhibited transcription was treated with 10 µg/mL Actinomycin-D. The control for background fluorescence did not contain 5EU (5EU-). After this incubation, cells where fixated with a 4% PFA solution in PBS for 15 minutes, followed by permeabilization with ice cold MeOH for 10 minutes at -20°C. Alexa-488 5-EU detection through Click-it chemistry was performed according to the manufacturer’s instructions (Click-iT™ HPG Alexa Fluor™ 488 Protein Synthesis Assay Kit; Invitrogen™ Cat#C10428). Nuclear staining was performed as described by the manufacturers of the kit. Imaging was performed in a CSU spinning disc confocal microscope (OLYMPUS with Andor iXon camera) with a 60x 1.4NA oil-immersion objective, for alexa-488 and nuclear staining signals. Imaging of a central z-slice per position was performed with a 488 nm laser at 35% power for 250 ms and a 405 nm laser at 20% power for 100 ms, with the gain set to 200.

#### RNA turnover assay

5×10^4^ mESCs where seeded per well in gelatinized µ-Slide Angiogenesis chip wells (ibidi Cat#81506) as described in the *experimental model and study participant details* section. 24h post-seeding, cells were exposed to 5EU containing medium (0.5mM) for 1h. A control with 5EU-medium was included. Following this incubation, the 5EU containing media was removed and substituted with regular mESC culture medium containing with CHX 250µM or corresponding volume of EtOH for 1 and 2 hours. Following this incubation, cells where fixated with a 4% PFA solution in PBS for 15 minutes, followed by permeabilization with ice cold MeOH for 10 minutes at -20°C. Alexa-488 5-EU detection through Click-it chemistry was performed according to the instructions of the providers of the detection kit (Click-iT™ HPG Alexa Fluor™ 488 Protein Synthesis Assay Kit; Invitrogen™ Cat#C10428). Nuclear staining was performed as described by the providers of the kit. Imaging was performed in a CSU spinning disc confocal microscope with a 60x 1.4NA oil-immersion objective, for alexa-488 and nuclear staining signals. Imaging of a central z-slice per position was performed with a 488 nm laser at 90% power for 500 ms and a 405 nm laser at 30% power for 200 ms, with the gain set to 200.

#### Poly-T Fluorescence in situ hybridization

5×10^4^ mESCs where seeded per well in gelatinized µ-Slide Angiogenesis chip wells (ibidi Cat#81506) after gelatinization. 24 hours post-seeding, cells were exposed to mESC culture medium containing 250 µM CHX, 10 µg/mL Actinomycin-D or the corresponding volume of EtOH for 1, 3 or 6 hours. After this incubation, cells where fixed with a 4% PFA solution in PBS and permeabilized with ice cold MeOH for 10 minutes at -20°C. Poly-A+ mRNA molecules were detected with Fluorescent *in situ* Hybridization (FISH) using 20-nucleotides-long Cy5.5 labeled oligo-T probes (Integrated DNA technologies; custom DNA oligos). In short, hybridization of oligo-T probes was performed overnight in hybridization buffer (HB) consisting of 1mL 20xSSC, 1mL 100% Formamide, 1g dextran sulfate and water up to 10 mL. Stock solution of probes (10 mM) was diluted 500:1 with HB. Hybridization was followed by one 30 minutes wash with washing buffer consisting of 5mL 20xSSC, 5mL 100% formamide and 40 mL water. The washing step was followed by DAPI staining, with 5 µg/mL DAPI in WB for 20 minutes. Imaging was performed in 2xSSC in a CSU spinning disc confocal microscope with a 60x 1.4NA oil-immersion objective, for Cy5.5 and DAPI signals. Imaging of a central z-slice per position was performed with a 640 nm laser at 40% power for 250 ms and a 405 nm laser at 20% power for 200 ms, with the gain set to 200.

#### RNA purification and bulk RNA sequencing

3×10^5^ mESCs were seeded per 9.6 cm^2^ well (Costar® 6-well Clear TC-treated Multiple Well Plate) after gelatinization. 24 hours post-seeding, cells were exposed to mESC culture medium with 250 µM CHX, for 3 or 6 hours. Control with corresponding volume of EtOH was treated for 9 hours. Samples where processed in duplicates. After this incubation, RNA extraction was performed with TRIzol^TM^ Reagent (Invitrogen^TM^ Cat#15596018) following the instructions of the manufacturer. Concentration and quality of the extracted RNA was assessed with Nanodrop One/Oneᶜ (ThermoFisher Scientific) and Bioanalyzer (Agilent Technologies). Frozen RNA samples were sent in dry ice to Singe Cell Discoveries (https://www.scdiscoveries.com/) for RNA sequencing.

#### Analysis of differential gene expression in bulk RNA sequencing after translation inhibition

Genome-wide read count values per sample were used as a starting point. DESeq2 package was used for analysis.^88^ Significant changes in mean were defined as adjusted p-value < 0.05.

#### Single-cell RNA sequencing of CHX treated cells

5×10^5^ mESCs were seeded in 9.6 cm^2^ wells (Costar® REF3516 plate). 24 hours post-seeding, the medium was changed to mESC culture medium with CHX 250 µM or equivalent volume of EtOH. Treatments with CHX proceeded for 3 or 6 hours. EtOH control proceeded for 6 hours. After treatment, cells were detached with 0.05% Trypsin in EDTA, centrifuged at 200×g for 3 minutes, resuspended in freezing medium consisting of 80% mESC culture medium, 10% DMSO and 10% extra FBS (Fisher Scientific Cat#A3840002), and immediately frozen at -80 °C. Frozen cells where shipped to the commercial company Single Cell Discoveries (https://www.scdiscoveries.com/) in dry ice for single-cell RNA sequencing with 10x Genomics technology. In brief, cells were thawed and encapsulated with cell-specific barcodes. Encapsulation was followed by reverse transcription and amplification through *in vitro* transcription. Finally, the prepared library was sequenced.

#### Bayesian analysis of differential gene expression in single-cell RNA sequencing data after CHX treatment

Genome-wide read count matrices for single cells were used as a starting point. For all three samples (i.e., EtOH control, 3h CHX and 6h CHX), cells were filtered based on their total read count and % of mitochondrial RNAs, utilizing Seurat V3.^89^ Thresholds were set based on the distributions of these parameters in the whole population of cells. Cells with total RNA count > 24000 and % of mitochondrial RNAs < 5% were kept for downstream analysis. Filtered Seurat objects were converted SingleCellExperiment objects^90^ and Marcov Chains Monte Carlo (MCMC) objects were obtained with BASiCS’ *BASiCS_MCMC* function.^37^ For this analysis, recommended settings for samples lacking spike-in RNAs were utilized (WithSpikes=FALSE, Regression=True, Prior=’log-normal’, Iterations=3000, Thin=10, Burn=750). MCMC allowed for Bayesian estimation of parameters describing the distribution of read counts in each population of cells. Differential expression testing for differential mean and differential variability (overdispersion, which is comparable to CV^2^) was performed utilizing BASiCS’ *BASiCS_TestDE* function. Significance is assessed through: (1) thresholds of 0.75 and 1.33 for the fold changes in mean or overdispersion, and; (2) a minimal value of 0.75 in posterior probability.

#### Liquid Chromatography coupled to tandem Mass Spectrometry

5×10^5^ mESCs were seeded in 9.6 cm^2^ wells (Costar® REF3516 plate). 24 hours post-seeding, the medium was changed to mESC culture medium with CHX 250 µM or equivalent volume of EtOH. Treatments with CHX proceeded for 3, 6 or 9 hours. EtOH control proceeded for 6 hours. 4 replicates for each treatment and control were performed. Lysis buffer was prepared by mixing 5.7 g of guanidine hydrochloride (Sigma-Aldrich Cat#G3272-500G), 1 mL HEPES 500 mM (Fisher Scientific Cat#1250979) and water up to 10 mL. After CHX treatment, medium was removed and 0.5 mL of lysis buffer was added to each well. Lysis buffer was pipetted up and down 3 times, making sure it covered the full well. Lysis buffer was transferred to a 1.5mL tube and stored at -80°C. Samples were sent in dry ice to the commercial company Biogenity (biogenity.com), that proceeded with the protein digestion, desalting of the samples and Liquid Chromatography coupled to tandem Mass Spectrometry analysis (TMT-labeled protein quantitation). 5323 proteins were detected in the 16 analyzed samples corresponding to 4 treatment conditions (control and CHX for 3, 6 and 9 hours) with 4 replicates. Each protein was tested for significant changes in mean for each CHX timepoint versus the control condition. Limma-moderated t-test was used for statistical comparisons, with significance set to p-value<0.05.

#### Regulator enrichment analysis

The groups of genes with significant overdispersion or mean changes in both 3h CHX sample and 6h CHX sample compared to the control sample were utilized as a starting point. A specific gene list was provided to ChEA3 gene set enrichment tool (https://maayanlab.cloud/chea3/ ; ^42^). Results for GTEx, ARCHS4, ReMap, ENCODE and Literature databases were downloaded. For each database, a list of Regulators whose defined targets are enriched in the input gene list were provided, with corresponding Odds Ratio and False discovery Rate (FDR). Regulators with FDR<0.05 were kept for downstream analysis. As the ChEA3 tool lacks an option to set a gene background for comparison, assessment of random enrichments was performed by providing the tool with 10 randomized gene list of size 2000 from the genes that do not changes their mean or overdispersion upon CHX treatment. Regulators enriched in more than one (i.e., 10%) randomized gene set were excluded from downstream analysis. Odds ratio was calculated as follows:

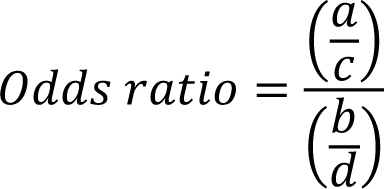

where *a* represents the genes from each list (i.e., our dataset) that are targets of a given regulator, *b* represents the genes from the background list (i.e., database) that are targets of a given regulator, *c* represents genes from each list (i.e., our dataset) that are NOT targets of the regulator, and *d* represents genes from the background list (i.e., database) that are NOT targets of the regulator.

#### Analysis of regulators target-set overlap and target exclusivity

The target set for each noise regulator as defined in the *Regulator enrichment analysis* section was utilized as starting point. Pairwise overlap between regulator target sets were computed as the number of shared targets and a graph representation of target overlap among regulator was obtained with the networknx package.^91^ Target exclusivity in regulator’s target sets was analyzed: For a given regulator (e.g. SON), a random target from its target set was sampled (e.g. *Yap1*), and the number of other regulators whose target set contained this same target (*Yap1*) was computed. This was repeated for all regulators in the specified category (e.g., noise enhancers). Two randomized controls were obtained: (1) a randomized intra-regulator control, by randomizing the target set of each regulator (not restricted to the genes subjected to a certain type of noise regulation) and repeating the previously described analysis; and (2) a global randomized control, by sampling 10 randomized regulator sets of the same size as the true regulator group (not restricted to a certain category of noise regulators) and repeating the previously described analysis for each random sample with the true targets of each.

#### Gene Ontology enrichment analysis

Gene ontology enrichment analysis of defined groups of genes or regulators versus specified background sets were performed utilizing WebGestalt. ^92^

#### siRNA mediated knock down (KD)

mESCs were seeded in experiment-specific number and culture format. 24 hours post-seeding cells were washed once with PBS and medium was changed to transfection medium. Transfection medium consisted of 70% mESC culture medium without penicillin nor streptomycin and 30% siRNA containing lipoplex solution. Lipoplex solution was obtained by combining 50% of 5% Lipofectamine^TM^ 2000 (Invitrogen^TM^ Cat#11668019) in Opti-MEM^TM^ (Gibco^TM^ Cat#31985062) with 50% of siRNA 1 mM in Opti-MEM^TM^, and incubating at room temperature for 15 minutes. Total volume of transfection medium was adjusted on an experiment-specific basis, depending on the cell number and culture format.

#### Validation of Son KD with immunostaining of intracellular protein and flow cytometry

mESCs were pelleted by centrifugation at 200×g for 3 minutes. Fixation was performed by resuspending the pellet in 0.5 mL of 4% paraformaldehyde in PBS and incubating for 15 minutes at room temperature. Following the incubation period, 5 mL of 0.2 mg/mL bovine serum albumin (BSA) were added and cells were pelleted by centrifugation at 300×g for 3.5 minutes. Permeabilization was performed by resuspending the pellet in 0.5 mL of ice cold MeOH and incubating for 10 minutes at -20°C. Following the incubation period, 5 mL of 0.2 mg/mL bovine serum albumin (BSA) were added and cells were pelleted by centrifugation at 300×g for 3.5 minutes. Blocking was performed by resuspending the pellet in 5% BSA in PBS and incubating for 1 hour at room temperature, while rotating. Following the blocking period, cells were split for isotype and protein-specific immunostaining, if required. Cells were pelleted by centrifugation at 300×g for 3.5 minutes. Primary or isotype antibody staining was performed by resuspending the cells in a 0.01 mg/mL dilution of isotype or anti-SON antibody (Invitrogen^TM^ Cat#PA5-115947) in 5% BSA in PBS, and incubated for 1 hour at room temperature, while rotating. Primary antibody staining was followed by 2 washes of 5 minutes with 0.1% Tween20 (PlusOne Cat#17-1316-01) in 0.2 mg/mL BSA in PBS. Staining with alexa488-labeled goat anti-rabbit secondary antibody (Invitrogen^TM^ Cat#A32731) was performed for all samples with concentration of 5 µg/mL, for 1 hour at room temperature, protected from light. Staining for secondary antibody was followed by one wash with 0.2 mg/mL BSA in PBS, after which cells where resuspended in PBS and analyzed in a BD FACSVerse flow cytometer.

#### Single-cell RNA sequencing of Son KD cells

1×10^6^ mESCs were seeded in two 25 cm^2^ flasks. 24 hours post-seeding, transfection with *Son* siRNA (ThermoFisher Scientific Cat#AM16704, ID:74755) or negative control siRNA (MISSION® siRNA Universal Negative Control #1) was performed as described in the *siRNA mediated knock down* section, with a total transfection medium volume of 7 mL. 24 hours post-transfection, the cells from each 25 cm^2^ flask were detached and split in two equal volumes. A sample of cell suspension was taken to assess viability as explained in the *Cell viability assay* section. Both equal volumes were pelleted by centrifugation at 200×g for 3 minutes. One was utilized to check the KD at the protein level, following the steps of the *Validation of KD with immunostaining of intracellular protein and flow cytometry* section. The second was resuspended in freezing medium, consisting of 80% mESC culture medium, 10% FBS (Fisher Scientific Cat#A3840002) and 10% DMSO, and immediately frozen at -80°C.

After validating the levels of KD as explained in the *Validation of KD with immunostaining of intracellular protein and flow cytometry* section, sample preparation for single-cell RNA sequencing with the frozen cells proceeded. Cells where thawed in a 37°C bath, pelleted and resuspended in PBS with BSA 0.04%. Cells were stained for viability with 7-AAD viability staining solution (ThermoFisher Scientific Cat#00-6993-50) for 5 minutes. As a decrease in viability in the *Son KD* sample was observed, sorting of viable cells prior to the single-cell RNA sequencing was performed. Viable cells were encapsulated together with cell-specific barcodes using 10x Genomics technology following the instructions of the manufacturer protocol (Chromium Next GEM Single Cell 3’ Reagent Kits v3.1). The sequencing of the prepared library was performed by the Radboud Sequencing Facility.

#### Bayesian analysis of differential gene expression in single-cell RNA sequencing data after Son KD

Genome-wide read count matrices for single cells was used as a starting point. For all two samples (negative control and *Son* KD), cells were filtered based on their total read count and % mitochondrial RNAs, utilizing Seurat V3^89^. Thresholds were set based on the distributions of these parameters in the whole population of cells. Cells with total RNA count > 4000 and % of mitochondrial RNAs < 5% were kept for downstream analysis. Filtered Seurat objects were converted to SingleCellExperiment objects^90^ and Marcov Chains Monte Carlo (MCMC) objects were obtained with BASiCS’ *BASiCS_MCMC* function ^37^. For this analysis, recommended settings for samples with spike in RNAs lacking were utilized (WithSpikes=FALSE, Regression=True, Prior=’log-normal’, Iterations=3000, Thin=10, Burn=750). MCMC allowed for Bayesian estimation of parameters describing the distribution of read counts in each population of cells. Differential expression testing for differential mean and differential variability (overdispersion) was performed utilizing BASiCS’ *BASiCS_TestDE* function. Significance is assessed through: (1) thresholds of 0.75 and 1.33 for the fold changes in mean or overdispersion, and; (2) a minimal value of 0.75 in posterior probability.

#### Quantification of cell cycle changes upon Son KD

***1×10***^6^ ***mESCs were seeded into two T-25 flasks. 24 hours post-seeding,*** transfection with *Son* siRNA (ThermoFisher Scientific Cat#AM16704, ID:74755) or negative control siRNA (MISSION® siRNA Universal Negative Control #1) was performed as described in the *siRNA mediated knock down* section, with a total transfection medium volume of 7 mL. To quantify cell cycle phases 24 hours post-KD, cells were incubated in media containing 5-Ethynyl uridine (EdU) 10 µM for 1 hour, followed by one wash with PBS, detachment with Trypsin 0.05% in EDTA for 3 minutes at 37; centrifugation at 200×g for 3 minutes and fixation by resuspending in 4% PFA for 15 minutes. Cells were washed with 0.2 mg/ml BSA in PBS and permeabilized with 100% MeOH at -20C. After permeabilizations cells were pelleted by centrifugation at 200×g for 3 minutes and EdU was detected following the instructions of the EdU proliferation kit (Abcam Cat#ab219801, iFluor 488). After detection cells were pelleted by centrifugation at 200×g for 3 minutes and stained with DAPI 1µg/ml in 0.2 mg/ml BSA in 1xPBS. Cells were pelleted by centrifugation at 200×g for 3 minutes and resuspended in PBS for flow cytometry analysis in a BD FACSVerse. Alexa-488 and DAPI signals were utilized to classify cells in G1 (low DAPI, low Alexa-488), G2 (high DAPI, low Alexa-488) or S (high Alexa-488, full range of DAPI).

#### Extrinsic contribution of cell cycle in single-cell RNA sequencing data

Each cell in the control and *Son* KD scRNA-seq datasets was attributed to a cell cycle phase (G1, S or G2). The differential gene expression analysis was repeated as explained in the *Bayesian analysis of differential gene expression in single-cell RNA sequencing data after Son KD* section, between cells attributed to the same cell cycle phase. After an equivalent filtering of the genome-wide read count matrices for single cells, the TriCycle package^93,94^ was utilized to attribute a θ value to each cell. In parallel, we calculated the z-scores of gene expression values and utilized the z-score values of Cyclins D and B (*Ccnd1* and *Ccnb1* genes), the two cycling cyclins in mESC cell cycle,^95,96^ to define the boundaries between cell cycle phases in the θ space.

#### Extrinsic contribution of heterogenous Son KD in single-cell RNA sequencing data

In order to estimate the impact of a heterogenous KD of *Son* on mRNA noise, clusters were obtained from both populations of cells sequenced in the scRNA-seq library upon *Son* KD. The RaceID package^97^ was used in order to obtain 10 clusters that can contain cells from both starting populations. In short, cell filtering was performed for control and *Son* KD populations as explained in *Bayesian analysis of differential gene expression in single-cell RNA sequencing data after Son KD* was performed, followed by the *pruneKnn* function of RaceID. In order to detect cells belonging to the population of *Son* KD cells that clustered together to control cells, the ratio of cells from each population in each of the 10 clusters was computed. Three types of clusters were detected: (1) predominantly control clusters; (2) predominantly *Son* KD clusters; and (3) mixed clusters. Cells originating from the *Son* KD population that were present in the predominantly control or mixed clusters were excluded, as these cells show evidences of poor or absent knock down. The analysis depicted in the section *Bayesian analysis of differential gene expression in single-cell RNA sequencing data after Son* KD was performed after excluding these cells, defined as “KD-escaping”.

#### Immunostaining of target proteins following Son KD

Target proteins and SON were co-stained directly in growing colonies when respective antibodies were compatible (i.e., for YAP1). When anti-target and anti-SON antibodies shared an isotype, the same population of transfected cells was split into different wells and attached with Cell-Tak^TM^ as explained in the *HPG incorporation assay* section (i.e., for ATF2). After the Cell-Tak^TM^ attachment (if required), cells were fixed with 4% paraformaldehyde in PBS for 15 minutes at room temperature. Fixation was followed by 3 washes with PBS. Permeabilization was performed with ice cold MeOH for 10 minutes at -20°C. Permeabilization was followed by 3 washes with PBS. Blocking was performed in 5% BSA in PBS, incubating for 1 hour at room temperature. Primary antibody staining was performed with a 0.01 mg/mL anti-SON antibody (Invitrogen^TM^ Cat#PA5-115947), 1 µg/mL anti-YAP1 antibody (Santa Cruz Biotechnology Cat#sc-101199) or 0.01 mg/mL anti-ATF2 antibody (Invitrogen^TM^ Cat#PA5-78831) in 5% BSA in PBS, and incubated for 1 hour at room temperature or overnight at 4 °C. Primary antibody staining was followed by 3 washes of 5 minutes with 0.1% Tween20 (PlusOne Cat#17-1316-01) in PBS. Staining with 5 µg/mL alexa488-labeled goat anti-rabbit secondary antibody (Invitrogen^TM^ Cat#A32731) and/or 2 µg/mL alexa647-labeled goat anti-mouse secondary antibody (Invitrogen^TM^ Cat#A21236) was performed for 1 hour at room temperature, protected from light. Staining for secondary antibody was followed by one wash with PBS, after which cells where stained with 5 µg/mL DAPI for 10 minutes in PBS and imaged in a CSU spinning disc confocal microscope with an oil 60x 1.4NA objective. Imaging of 0.5 µM spaced z-slices spanning the entire thickness of cells per position was performed with a 488 nm laser at 30% power for 100 ms or 50% power for 120 ms, a 405 nm laser at 30% power for 50 or 120 ms, and a 641 nm laser at 50% power for 300 ms, with the gain set to 200. Cells were segmented with Cellpose’s AI-based pretrained models.^98,99^ Protein staining for SON or ATF2 were utilized as the input for cell segmentation. Cell masks provided by Cellpose were utilized in ImageJ to obtain single-cell measurements of average protein staining signal in the corresponding SON, ATF2 or YAP1 channels. For mean and noise analysis, extrinsic noise filtering was applied where cells were filtered based on a range of areas (1000-3000 pixel^2^)^18^. Noise was quantified through the CV^2^ (σ^2^/µ^2^) and the Fano factor (σ^2^/µ). In each experiment, in order to exclude the effect of outliers, randomized groups of 100 to 200 cells from the control and KD samples were sampled and mean and noise changes were quantified per subsampled population. Significance of changes in noise where computed for the average Fano factor values. Significance was computed with independent sample t-test, with a significance threshold of p-value<0.05.

#### Long-read RNA sequencing of Son KD cells

1×10^6^ mESCs were seeded in two 25 cm^2^ flasks. 24 hours post-seeding, transfection with *Son* siRNA (ThermoFisher Scientific Cat#AM16704, ID:74755) or negative control siRNA (MISSION® siRNA Universal Negative Control #1) was performed as described in the *siRNA mediated knock down* section, with a total transfection medium volume of 7 mL. 24 hours post-transfection, the cells from each 25 cm2 flask were detached and split in two equal volumes. Both equal volumes were pelleted by centrifugation at 200×g for 3 minutes. One was utilized to check the KD at the protein level, following the steps of the *Validation of KD with immunostaining of intracellular protein and flow cytometry* section. RNA was extracted from the second volume of cells with TRIzol^TM^ Reagent (Invitrogen^TM^ Cat#15596018) following the instructions of the manufacturer. Concentration and quality of the extracted RNA was assessed with Nanodrop One/Oneᶜ (ThermoFisher Scientific) and Bioanalyzer (Agilent Technologies). Library preparation for long-read RNA sequencing was performed with the cDNA-PCR Sequencing Kit (SQK-PCS111) in one R9.4.1 Flow Cell per sample, in a MinION device. Library preparation was performed as described in the protocol provided by the manufacturer (cDNA-PCR Sequencing), starting from 200 ng of total RNA. Validation of library quality was performed with Qubit and Bioanalyzer (Agilent Technologies), with a DNA 12000 chip. Basecalling of registered electric signals was performed in parallel with sequencing by the software provided by the manufacturer.

#### Analysis of long-read RNA sequencing upon Son KD

Alignment of reads to a reference mouse transcriptome was performed with minimap2 ^100^. The transcriptome of reference is GRCm39, downloaded from Ensembl database (^101^ https://ftp.ensembl.org/pub/release-110/fasta/mus_musculus/). Counts per transcript variant were obtained utilizing Samtools ^102^. Variant count matrices in the control and *Son* KD samples were normalized to transcripts per million (TPM) to account for different library sizes. Transcript variants were filtered based on their overall TPM (TPM_control_ + TPM_KD_). Specifically, variants with log_2_(overall TPM) < -2 were excluded from downstream analysis. Usage (U) of each variant was quantified as the percentage of all TPMs for the corresponding gene that are associated with each specific variant, as defined by the following equation:

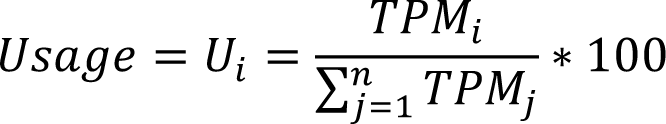

where i indicates one specific transcript variant, and j indicates all transcript variants for the same gene as i, from 1 to n. Directional and absolute usage changes were calculated as follows:

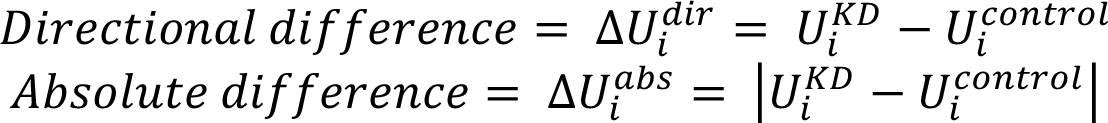

Transcript and gene properties corresponding to each transcript variant were downloaded from Ensembl database through BioMart ^103,104^. These properties include transcript biotype as defined in the Ensembl database. Absolute usage differences were combined in cumulative absolute usage differences, by adding up all the absolute usage differences for variants belonging to the same biotype, as defined by the following equation:

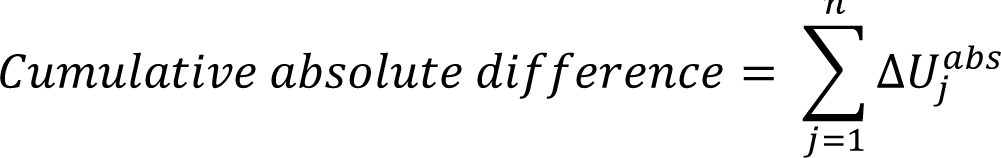

where j indicates all transcript variants for the same gene that belong to the same biotype, from 1 to n. Genes were classified in five categories based on the results of differential mean and noise as described in *Bayesian analysis of differential gene expression in single-cell RNA sequencing data after Son KD*. These five categories were: mean enhanced, mean repressed, noise enhanced, noise repressed and unchanged. For each category and variant biotype, the average and standard error of the mean of cumulative absolute usage differences were computed. For each category and variant biotype, the average and standard error of the mean of directional usage differences were also computed, when considering only the most common transcript variant for a gene in the control condition. Significance of differences between each category (mean enhanced, mean repressed, noise enhanced and noise repressed) and the control unchanged group were computed with independent samples t-test provided that the variances of both samples showed a difference <4-folds, which is, in general terms, the case for all comparisons. Significance threshold is set to p-value < 0.05. In Figure S6D, boxplots represent the 25^th^, 50^th^ and 75^th^ percentiles of the distribution of means obtained through a subsampling strategy where 20 randomized groups of 500 genes are considered (independently of their mean/noise change category).

Inverse Simpson’s index was calculated, for each specific gene, utilizing the following equation:

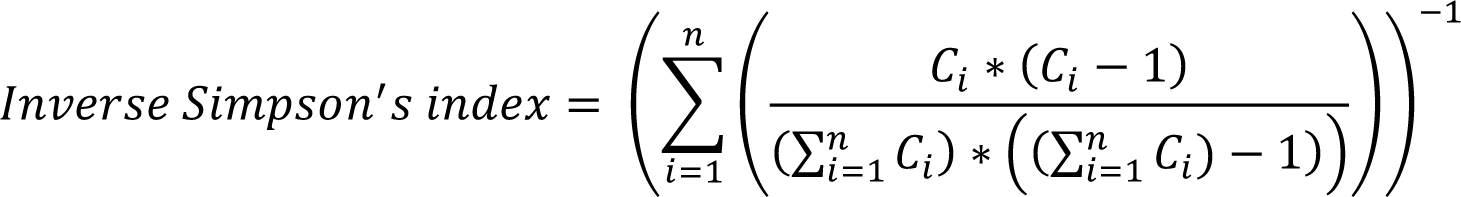

where C_i_ is the number of reads counted for the transcript isoform I, and n is the number of isoforms of that given gene.

#### Estimation of bursting kinetics with nnRNA

The single-cell gene expression matrices for control and *Son* KD samples after filtering as in the *Bayesian analysis of differential gene expression in single-cell RNA sequencing data after Son KD* section were utilized as the starting point. In order to increase the accuracy of estimated parameters, a second filtering step for genes with mean expression >=0.1 was included. Estimation of bursting kinetics was performed with the function^66^:

> *nnRNA_wrapper.nnRNA_wrapper(scRNA=scRNA_input,Gname=Gname_input,file_name=fi le_name,store_path=store_path,allele_double=True,repeats=1,threshold_gc=[5000,2,100], prior=“Fano”,inputbeta_vec=inputbeta)*

with inputbeta = sim.empirBETA(scRNA_input,meanBeta=0.06) and Gname_input=None. Note that *allele_double=True* was utilized as the data lacked allele specificity. Columns 0, 1, 2 and 3 of the output table correspond to the log_10_ of the gene-specific estimations of k_on_, k_off_, k_syn_ and k_syn_/k_off_ respectively, with each row corresponding to a gene in the order of the input matrix.

Average basal values for these parameters (control sample) and changes in these parameters between the *Son* KD and the control samples were assessed in 10 randomized subsets of 10% of genes included in each category of mean/noise modulation, and error of mean changes was quantified as the standard error of the mean in this subsampling process.

#### Total RNA sequencing and analysis of Splicing efficiency

The same isolated RNA obtained from the *Long-read RNA sequencing of Son KD cells* section was utilized as one of the two replicates for this sequencing experiment. A second biological replicate obtained with the same procedure (that was not subjected to Long-Read RNA sequencing) was also obtained following the same procedure. Total RNA was sent in dry ice to the commercial company GenomeScan (www.genomescan.nl) for total RNA sequencing, where quality control of the samples was performed for a second time. rRNA was depleted from the sample and the remaining RNA was fragmented and utilized in the library preparation for sequencing with NovaSeq6000. Produced paired-end reads were aligned with STAR2.7.11a^105^ against the *Mus musculus* genome of reference GRCm39, downloaded from Ensembl. Output .bam files were sorted with Samtools ^102^. These sorted alignment files together as the indexed .gtf file of *Mus musculus* GRCm39 were utilized as the input for the analysis of splicing efficiency with Splice-q ^62^. The file SPLICE-q.py was downloaded from the author’s GitHub repository (https://github.com/vrmelo/SPLICE-q) and utilized as indicated by the developers.

#### Speckle disruption analysis upon tubercidin treatment

To visualize nuclear speckles, SON was stained and imaged following as defined in *Immunostaining of target proteins following Son KD.* Nuclei were segmented with Cellpose’s AI-based pretrained models,^98,99^ utilizing DAPI channel as input. A threshold of 10 000 a.u. was applied to the SON immunofluorescence images to obtain a binary mask of the speckles. In order to detect areas of high intensity, the threshold was established by quantifying the average intensities of randomized regions (of ±1.17 µm^2^) within segmented nuclei. The ±95^th^ percentile of this randomized signal was set as the threshold. Nuclear masks provided by Cellpose were utilized in ImageJ to obtain single-nuclei measurements of total SON intensity per cell as well as number, size, and intensity of SON speckles according to the previously described masks and within an area range of 0.23-2.33 µm^2^.

#### Flow cytometry analysis of NANOG expression upon Tubercidin treatment

5×105 NANOG-GFP mESCs were seeded in gelatin coated in 9.6 cm^2^ wells (Costar® REF3516 plate). 24 hours post-seeding, Tubercidin (Fisher Scientific Cat# HY-100126_5mg) 1 µM or 10 µM, or the corresponding volume of DMSO was introduced in the mESC culture medium. Flow cytometry analysis of GFP signal was performed with a BD FACSCalibur Flow cytometer 6 hours post-treatment. In short, GFP expression and noise was calculated after extrinsic noise filtering (Figure S6A-B) was performed.^18^ Data analysis was performed with BD FlowJo.

#### Single-cell RNA sequencing of Tubercidin-treated cells

5×10^5^ mESCs were seeded in 9.6 cm^2^ wells (Costar® REF3516 plate). 24 hours post-seeding, the medium was changed to mESC culture medium with Tubercidin (Fisher Scientific Cat# HY-100126_5mg) 10 µM or equivalent volume of DMSO. Treatments proceeded for 3 hours. Cells where then detached, pelleted, resuspended in PBS with BSA 0.04% and brought fresh to the encapsulation step. Cells were encapsulated together with cell-specific barcodes using 10x Genomics technology following the instructions of the manufacturer protocol (Chromium Next GEM Single Cell 3’ Reagent Kits v3.1). The sequencing of the prepared library was performed by the Radboud Sequencing Facility.

#### Bayesian analysis of differential gene expression in single-cell RNA sequencing data after Tubercidin treatment

Genome-wide read count matrices for single cells were used as a starting point. For all two samples (DMSO control, 3h TUB), cells were filtered based on their total read count and % of mitochondrial RNAs, utilizing Seurat V3.^89^ Thresholds were set based on the distributions of these parameters in the whole population of cells. Cells with total RNA count > 4000 and % of mitochondrial RNAs < 5% were kept for downstream analysis. Filtered Seurat objects were converted SingleCellExperiment objects^90^ and Marcov Chains Monte Carlo (MCMC) objects were obtained with BASiCS’ BASiCS_MCMC function (Vallejos et al., 2015). For this analysis, recommended settings for samples lacking spike-in RNAs were utilized (WithSpikes=FALSE, Regression=True, Prior=’log-normal’, Iterations=3000, Thin=10, Burn=750). MCMC allowed for Bayesian estimation of parameters describing the distribution of read counts in each population of cells. Differential expression testing for differential mean and differential variability (overdispersion, which is comparable to CV2) was performed utilizing BASiCS’ BASiCS_TestDE function. Significance is assessed through: (1) thresholds of 0.75 and 1.33 for the fold changes in mean or overdispersion, and; (2) a minimal value of 0.75 in posterior probability.

#### Analysis of transcript intronic properties and association with nuclear speckles in relation to noise regulation

The exon distribution of genes was retrieved from Ensembl database with BioMart ^103,104^. For each gene and isoform intron lengths were computed as the difference between the starting genomic location of the exon n+1 and the end location of the exon n. From this information, values of intron lengths, number of introns, average intron length, total intronic length, CV of intron lengths, and lengths of the shorter and longer introns were computed for each isoform. Genes were classified according to a combined category in terms of noise or mean regulation as defined by both *Son* KD and Tubercidin datasets. 10 random samples of 100 genes were drafted from each combined category and the average behavior in regards to the previously mentioned intronic properties were calculated.

Regarding the association to nuclear speckles, a total of 4552 genes were associated to the active compartment (associated with nuclear speckles, 3119 genes) or nucleolar compartment (anti-associated with nuclear speckles, 1433 genes).^74^ From these groups of genes, 1580 and 345 respectively were present in the *Son* KD scRNA-seq dataset, and 1505 and 343 in the Tubercidin treatment dataset. Only genes present in either of the two categories were included in this analysis. 20 randomized samples of 50% of genes where obtained from each group of noise regulated genes as identified through *Son* KD or Tubercidin treatment, and the Active/nucleolar ratio was defined as the ratio between the count of genes associated with each compartment, for each subset of genes.

#### mESC differentiation efficiency assay

8.4×10^3^ mESCs were seeded per well in gelatin coated Costar® 24-well Clear TC-treated Multiple Well Plate (Cat#CL23526). 24 hours post-seeding, KD of SON was induced as described in the *siRNA mediated knock down (KD)* section, with *Son* siRNA (ThermoFisher Scientific Cat#AM16704, ID:74755) or negative control siRNA (MISSION® siRNA Universal Negative Control #1). 24 hours post-transfection, transfection medium was removed and cells were grown in differentiation medium. Differentiation medium consisted of 50% Neurobasal medium (Gibco^TM^ Cat#21103049) and 50% DMEM F12 (Capricorn Scientific Cat#DMEM-12-HXA), supplemented with L-glutamine (Thermo Scientific™ Cat#25030024), mercaptoethanol (Fisher Scientific Cat#11528926), N2 (Gibco^TM^ Cat#17502048) and B27 (Gibco^TM^ Cat#17504044). In the experiments involving external noise enhancement, 4 µM IdU or corresponding volume of DMSO was added to differentiation medium. Differentiation proceeded for 96 hours. Differentiation efficiency was assessed by the immunostaining of membrane markers CD24 in live cells with Alexa Fluor® 647 anti-mouse CD24 Antibody (BioLegend Cat#101818). Isotype control Alexa Fluor® 647 Rat IgG2b, κ Isotype control (BioLegend Cat#400626) was used to gate CD24+ cells. The assay was performed in 6 biological replicates, and significant changes between conditions were assessed with independent samples t-test with a significance threshold of p-value<0.05.

#### mESC fate choice assay

4×10^4^ NANOG-GFP mESCs were seeded per well in gelatin coated Costar® 24-well Clear TC-treated Multiple Well Plate (Cat#CL23526). 24 hours post-seeding, KD of SON was induced as described in the *siRNA mediated knock down (KD)* section, with *Son* siRNA (ThermoFisher Scientific Cat#AM16704, ID:74755) or negative control siRNA (MISSION® siRNA Universal Negative Control #1). 24 hours post-transfection, transfection medium was removed and cells were grown in differentiation medium. Differentiation medium consisted of 50% Neurobasal medium (Gibco^TM^ Cat#21103049) and 50% DMEM F12 (Capricorn Scientific Cat#DMEM-12-HXA), supplemented with L-glutamine (Thermo Scientific™ Cat#25030024), mercaptoethanol (Fisher Scientific Cat#11528926), N2 (Gibco^TM^ Cat#17502048) and B27 (Gibco^TM^ Cat#17504044). Retinoic acid (Sigma-Aldrich Cat#R2625) was added to the differentiation medium to a concentration of 0.25 µM. Differentiation proceeded for 96 hours. Fate choice during differentiation was assessed by the immunostaining of membrane markers CD24 in live cells with Alexa Fluor® 647 anti-mouse CD24 Antibody (BioLegend Cat#101818), with Isotype control Alexa Fluor® 647 Rat IgG2b, κ Isotype control (BioLegend Cat#400626) used to gate CD24+ cells; and the immunostaining of membrane markers CD140a in live cells with Brilliant Violet^TM^ 421 anti-mouse CD140a Antibody (BioLegend Cat#135923), with Isotype control Brilliant Violet^TM^ 421 Rat IgG2b, κ Isotype control (BioLegend Cat#400535) used to gate CD140a+ cells. Endogenous NANOG-GFP signal was also measured. The assay was performed in 3 biological replicates at 5 timepoints during differentiation (0, 24, 48, 72 and 96 hours).

#### Repopulation of NANOG-GFP states

5×10^5^ NANOG-GFP mESCs were seeded in gelatin coated in 9.6 cm2 wells (CostarR REF3516plate). 24 hours post-seeding, KD of SON was induced as described in the siRNA mediated knock down (KD) section, with Son siRNA (ThermoFisher Scientific Cat#AM16704, ID:74755) or negative control siRNA (MISSIONR siRNA Universal Negative Control #1). 24 hours post-transfection cells were washed once with PBS, detached with Accutase^TM^ (STEMCELL^TM^ Technologies Cat#07920) for 5 minutes at 37°C, pelleted by centrifugation at 200 rcf for 3 minutes and resuspended in 1xPBS. Utilizing a cell sorter of the model BD FACSAria cells were gated based on FCS-A/SSC-A and FSC-H/FSC-A gates (healthy single cells) and then sorted based on their GFP signal, as a proxy for NANOG. To define the two sorted populations, the GFP distribution was split in two (GFP-low and GFP-high) containing approximately 50% of cells. The ratio between the sorted populations (in terms of GFP median) was measured to be around 2-fold. Sorted cells were deposited in mESC culture medium (serum/LIF) without penicillin nor streptomycin. Cell count and viability was measured in each sorted population as explained in the Cell viability assay section. 4 ×10^4^ cells were seeded per gelatin coated well of a Costar® 48-well Clear TC-treated Multiple Well Plate (Cat#CLS3548). 24 or 48h post-seeding, cells were washed with PBS and detached with Accutase^TM^ (STEMCELL^TM^ Technologies Cat#07920) for 5 minutes at 37°C for 5 minutes at 37°C, followed by flow cytometry analysis of GFP signal with a BD FACSVerse Flow cytometer. If indicated, IdU was introduced in the growing media after 24h of post-sorting growth. Repopulation was computed as described by the following equation for 10 randomized samples of 1000 cells from each sorted population (high and low), in 2 biological replicates.

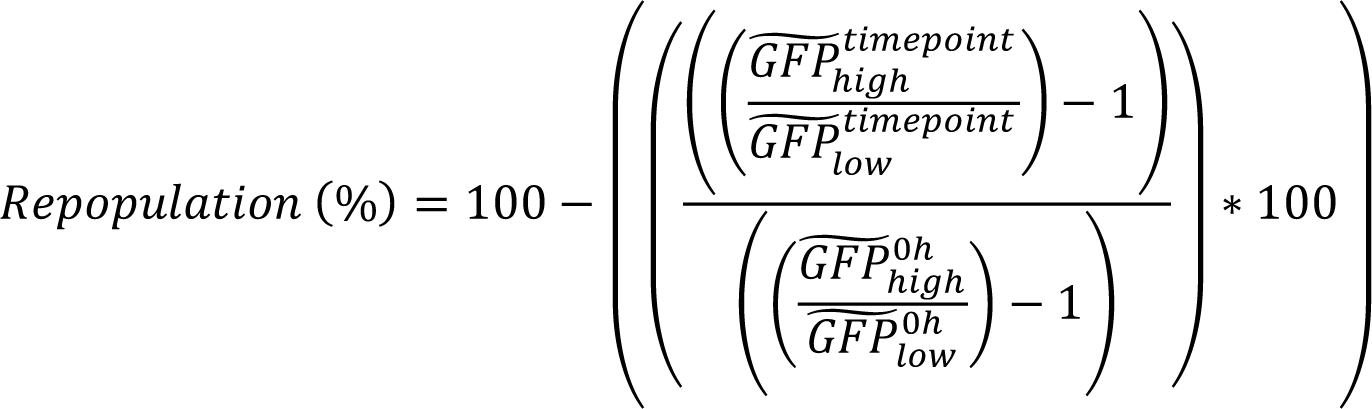

#### CD9 expression assay

4×10^4^ mESCs were seeded per gelatin coated well of a Costar® 24-well Clear TC-treated Multiple Well Plate (Cat#CL23526). 24 hours post-seeding, KD of SON was induced as described in the *siRNA mediated knock down (KD)* section, with *Son* siRNA (ThermoFisher Scientific Cat#AM16704, ID:74755) or negative control siRNA (MISSION® siRNA Universal Negative Control #1). 24 hours post-transfection, transfection medium was removed and cells were grown in mESC culture medium as described in the *Culture of mouse embryonic stem cells* section, but in the absence of Penicillin and Streptomycin. At this point, 4 µM 5-Iodo-2’-deoxyuridine (IdU; Merck Cat#I7125) or corresponding volume of DMSO was added to the mESC culture medium. 24 hours later, CD9 membrane marker was stained in live cells, with FITC anti-mouse CD9 Antibody (BioLegend Cat#124808) or corresponding FITC Rat IgG2a, κ Isotype control Antibody (BioLegend Cat#400506) for 30 minutes at 37°C in the same mESC culture medium as previously described. Following the staining, cells were detached with Accutase^TM^ (STEMCELL^TM^ Technologies Cat#07920) for 5 minutes at 37°C, colonies were mechanically disrupted by repeated pipetting and cells were analyzed in a BD FACSVerse flow cytometer. Data analysis was performed with BD FlowJo. The assay was performed in 6 biological replicates, and significant changes between conditions were assessed with independent samples t-test with a significance threshold of p-value<0.05.

#### Analysis of SON expression during *mESC differentiation*

4×10^3^ mESCs were seeded per well in gelatin coated µ-Slide 18 well (ibidi Cat#81816). 24 hours post-seeding, regular growth medium was removed and cells were grown in differentiation medium. Differentiation medium consisted of 50% Neurobasal medium (Gibco^TM^ Cat#21103049) and 50% DMEM F12 (Capricorn Scientific Cat#DMEM-12-HXA), supplemented with L-glutamine (Thermo Scientific™ Cat#25030024), mercaptoethanol (Fisher Scientific Cat#11528926), N2 (Gibco^TM^ Cat#17502048) and B27 (Gibco^TM^ Cat#17504044). At defined differentiation timepoints of 0h, 48h and 96h, cells were fixed with 4% paraformaldehyde in PBS for 15 minutes at room temperature. Fixation was followed by 3 washes with PBS. Permeabilization was performed with ice cold MeOH for 10 minutes at - 20°C. Permeabilization was followed by 3 washes with PBS. Blocking was performed in 5% BSA in PBS, incubating for 1 hour at room temperature. Primary antibody staining was performed with a 0.01 mg/mL anti-SON antibody (Invitrogen^TM^ Cat#PA5-115947 in 5% BSA in PBS, and incubated overnight at 4 °C. Primary antibody staining was followed by 3 washes of 5 minutes with 0.1% Tween20 (PlusOne Cat#17-1316-01) in PBS. Staining with 5 µg/mL alexa488-labeled goat anti-rabbit secondary antibody (Invitrogen^TM^ Cat#A32731) was performed for 1 hour at room temperature, protected from light. Staining for secondary antibody was followed by one wash with PBS, after which cells where stained with 5 µg/mL DAPI for 10 minutes in PBS and imaged in a CSU spinning disc confocal microscope with an oil 60x 1.4NA objective. Imaging of 0.5 µM spaced z-slices spanning the entire thickness of the cells per position was performed with a 488 nm laser at 12% power for 50 ms, a 405 nm laser at 40% power for 50 ms, with the gain set to 172. Cells were segmented with Cellpose’s AI-based pretrained models ^98,99^. The maximum z-projection of DAPI channel was utilized as the input for cell segmentation. Cell masks provided by Cellpose were utilized in ImageJ to obtain single-cell measurements of average SON staining signal in maximum z-projections of corresponding channel.

### QUANTIFIATION AND STATISTICAL ANALYSYS

The information regarding replicates, statistical analysis and significance is included in each specific *method details* section and in corresponding figures legends, when required. Most statistical analysis have been performed in Python 3^106^, with SciPy^107^ as the main package for statistical analysis. Specific R packages (BASiCs^37^) or web services (ChEA3^42^, WebGestalt^92^) have been used in specific analysis steps. Graphical representations of statistical analysis are performed in Python 3 with matplotlib^108^ and seaborn^109^.

## SUPPLEMENTARY TABLES

**Table S1. Related to Figure S3.** Gene Ontology enrichment analysis of enriched regulators for noise enhanced, mean enhanced and mean repressed genes upon CHX treatment.

**Table S2. Related to** Figure 7. Gene Ontology enrichment analysis of targets for mean/noise regulation in *Son* KD.

## SUPPLEMENTARY DATA

**DataS1.** Proteomics dataset obtained from protein extracts upon translation inhibition with cycloheximide for 3, 6 or 9h, or respective control. Data for peptides.

**DataS2.** Proteomics dataset obtained from protein extracts upon translation inhibition with cycloheximide for 3, 6 or 9h, or respective control. Data for proteins.

**DataS3.** Proteomics dataset obtained from protein extracts upon translation inhibition for 3,6 or 9h, or respective control. Key to relate sample numbers to treatment timepoints in DataS1 and DataS2.

