## Supplementary Figures 1-7 for "Multimodal screen reveals noise regulatory proteins"

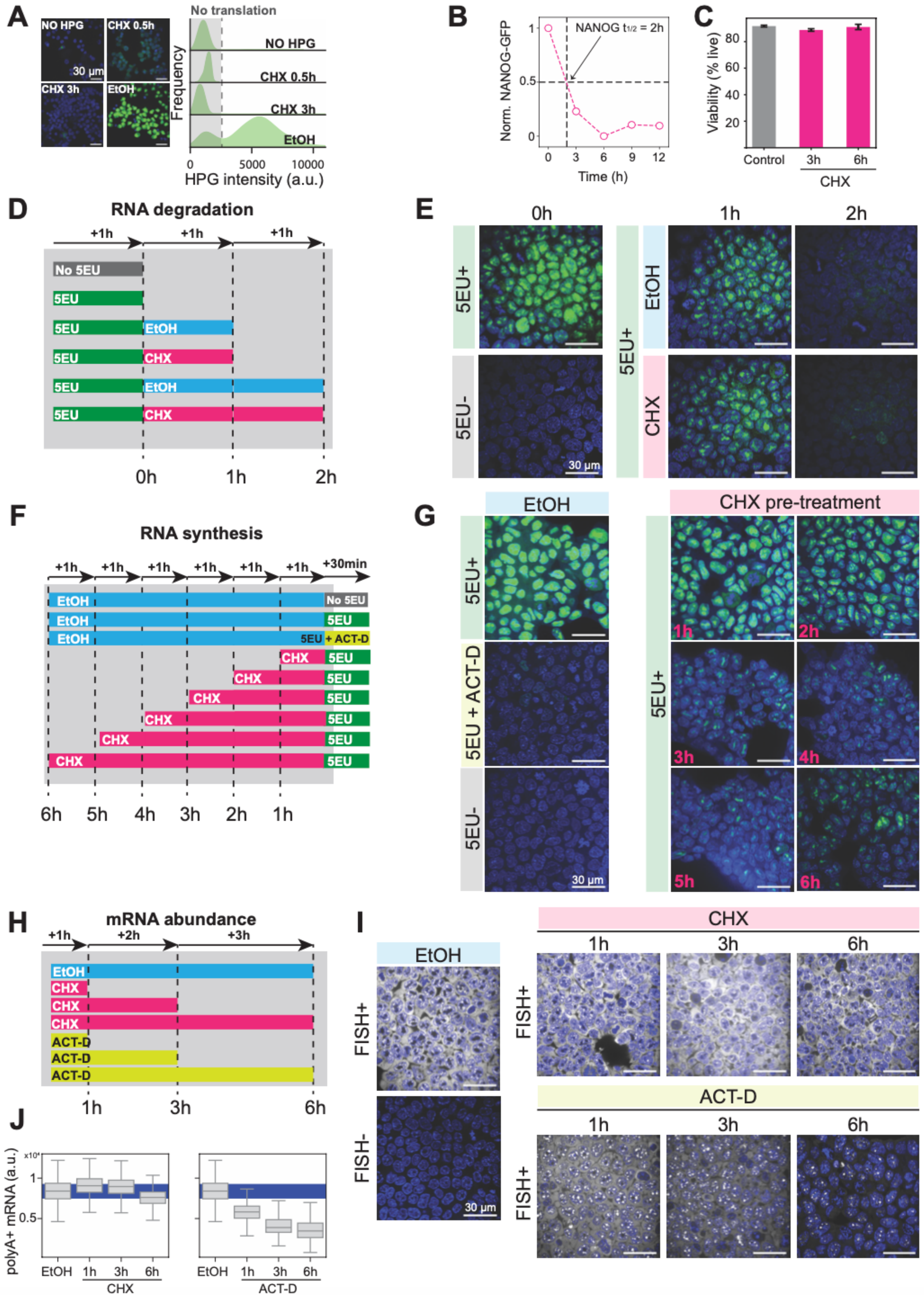


**Figure S1. CHX treatment alters total RNA synthesis, while total RNA degradation and mRNA levels remain unaffected, Related to Figure 1.**

**(A-C)** Inhibition of translation with cycloheximide. (A) Representative microscopy images and distribution of incorporated l-homopropargylglycine (HPG) in single mESCs treated with 250 μM cycloheximide (CHX) for 0.5-3 hours or EtOH (control sample). Shaded area represents the background fluorescence measured in the No HPG control. (B) Normalized expression of NANOG-GFP in NANOG-GFP mESCs over time following treatment with 250 μM CHX with respect to EtOH (control sample). (C) mESC viability after treatment with 250 μM CHX for 3-6 hours or EtOH (control sample). Error bars represent standard error of the mean.

**(D)** Schematic of the 5EU incorporation assay for nascent total RNA transcription.

**(E)** Representative microscopy images of the 5EU incorporation assay for nascent total RNA transcription.

**(F)** Schematic of the 5EU decay assay for total RNA turnover.

**(G)** Representative microscopy images of the 5EU decay assay for total RNA turnover.

**(H)** Schematic of the control sample (EtOH), CHX and ACT-D treatments for polyA+ FISH.

**(I)** Representative microscopy images of polyA+ FISH upon CHX or ACT-D treatments.

**(J)** Box plots of single-cell polyA+ FISH signal in the control sample (EtOH), CHX and ACT-D treatment. Blue shaded area represents the range of 90 to 110% change of the EtOH control.

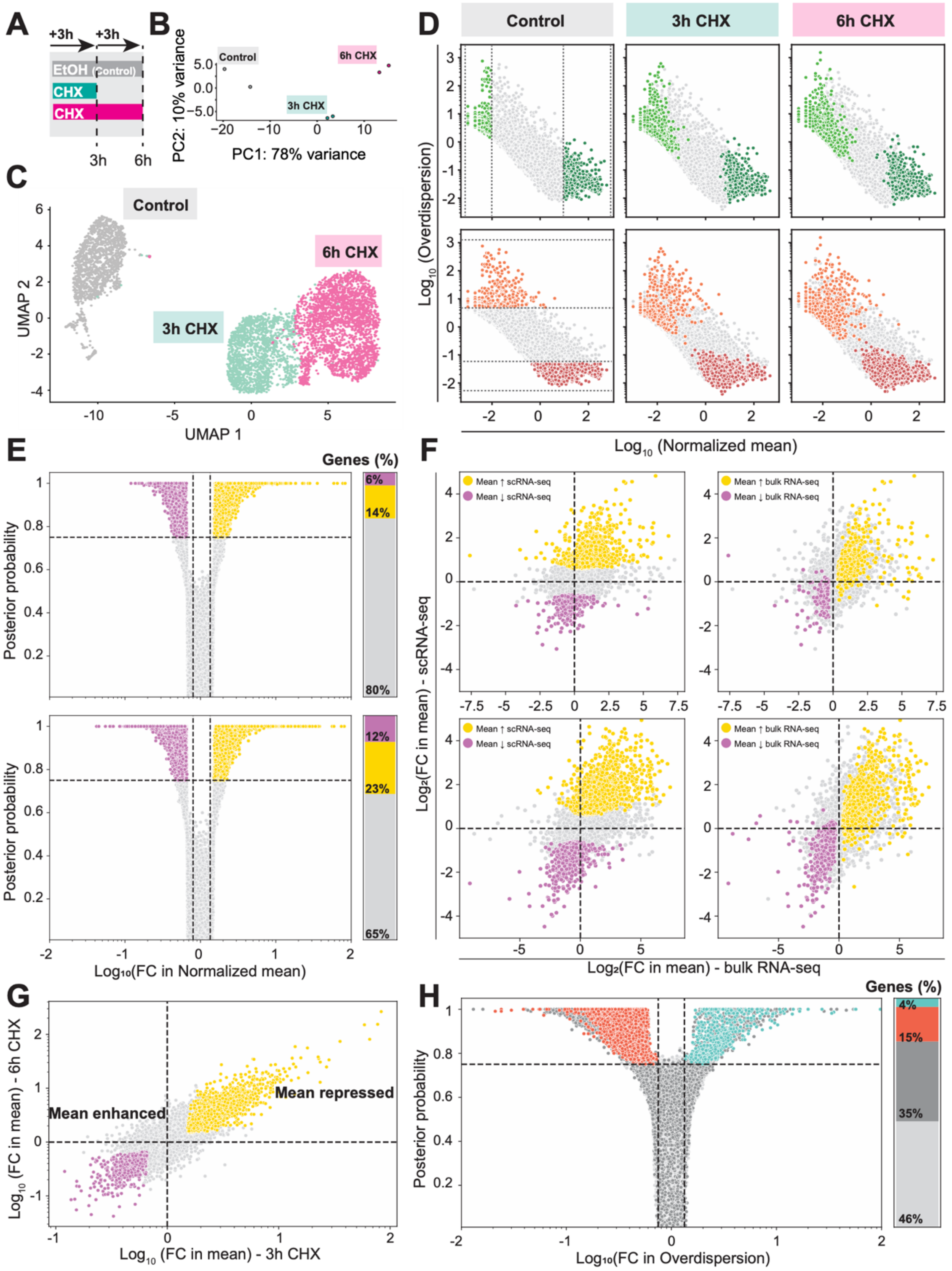


**Figure S2. Extended analysis of scRNA-seq upon CHX treatment, Related to Figure 1.**

**(A)** Schematic of CHX treatment.

**(B)** Principal component analysis of EtOH control and 3- and 6-hour CHX treatment, according to the schematic in (A), after bulk RNA-seq, with two replicates per sample type.

**(C)** UMAP dimensionality reduction showing distinct clustering of the control sample and 3- and 6-hour CHX treatment (pink, turquoise and grey respectively), after scRNA-seq.

**(D)** Normalized mean expression versus overdispersion from scRNA-seq of control mESCs and treated with 250 μM of CHX for 3 and 6 hours (1462, 1560, 2395 cells respectively) with highlighted genes displaying a high/low range in mean expression (top, light/dark green) and overdispersion (bottom, red/orange) in the control.

**(E)** Differential expression analysis, for differential mean with BASiCs of 250 μM CHX treatment for 3 (top) and 6-hours (bottom) versus control sample. Volcano plot represents the fold change versus the certainty (Posterior probability) that this change is significant (significance thresholds are depicted with dashed lines). Stacked bar plot represents the % genes that display significant changes in mean (3398 in 3h treatment; 5809 in 6h treatment).

**(F)** Comparison between mean expression changes measured with bulk RNA-seq and scRNA-seq, for 3h CHX vs control sample (top), and 6h CHX vs control sample (bottom). Significant mean changers are highlighted in yellow or violet for scRNA-seq (left) or bulk RNA-seq (right).

**(G)** Fold changes in mean expression of the 3-hour CHX sample compared to the control sample versus the 6-hour CHX sample compared to the control sample. Genes showing a consistent significant change in mean expression are highlighted in yellow (2179 mean-repressed genes) and violet (742 mean-enhanced genes).

**(H)** Differential expression analysis, for differential overdispersion with BASiCs of 250 μM CHX treatment for 6-hours (bottom) versus the control sample. Volcano plot represents the fold change versus the certainty (Posterior probability) that this change is significant (significance thresholds are depicted with dashed lines). Stacked bar plot represents the % genes that display significant changes in noise (overdispersion; orange: 2512 noise-enhanced genes; turquoise: 613 noise-repressed genes; dark grey: 5809 genes excluded because they display differential mean expression).

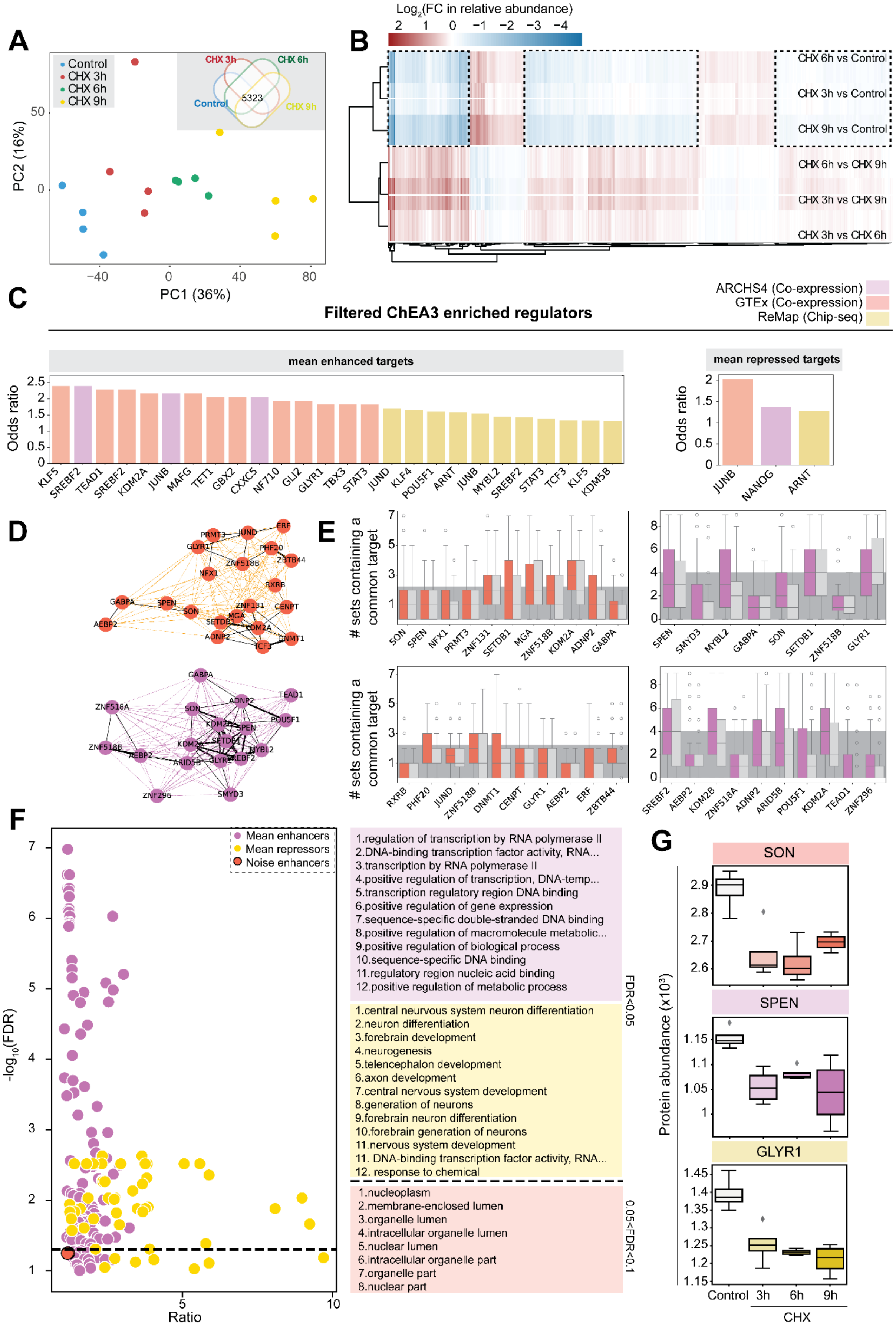


**Figure S3. Multimodal screen identifies mean regulators, Related to Figure 2.**

**(A-B)** Detailed analysis of proteomics data. (A) Principal component analysis of 16 samples included in LC-MC/MC analysis. Venn diagram of number of proteins detected in all samples measured by proteomics. 5323 common proteins were detected with confidence in all samples (inset). (B) Heatmap of 5323 proteins detected in 6 comparisons, showing expression levels changes (Log2(Fold Change in relative abundance)). Both proteins and comparisons have been ordered according to hierarchical clustering. Clusters marked with dashed lines represent short-lived proteins of interest.
**(C-G)** Extended regulator enrichment analysis. (C) Bar plot of significant candidate regulators identified through regulator enrichment analysis of mean-repressed targets (left) and mean-enhanced targets (right). Regulators are ranked according to their Odds ratio. The color of the bar represents the database of origin of the regulator and its respective target set. (D) Network graph of target-set overlaps for regulators identified as noise enhancers in Figure 2C. Nodes represents the regulators from GTEX (top; orange) or ARCHS4 (bottom; violet) databases. Edge color and thickness represents the overlap between the target sets of the two connected regulators: black thick edge corresponds to >25 shared targets, black thin edge correspond to <25 and >10 shared targets, and colored edge represents <10 shared targets. (E) Analysis of target exclusivity for regulators identified as noise enhances from GTEX (top; orange) or ARCHS4 database (bottom; violet). Light grey boxes correspond to equivalent analyses with randomized target sets. Dark grey shaded delimits the 5 to 95% percentiles of an equivalent analysis from a randomized set of regulators. (F) Gene set enrichment analysis for gene ontology terms in groups of enriched regulators from mean enhancers (from Figure 4D, left; in violet), repressors (from Figure S4D, right; in yellow), and noise (overdispersion) enhancers (from Figure 3C; in orange), without filtering regulators based on their half-life upon CHX treatment, compared to a background of all regulators included in Chea3. Data is represented as a scatter plot (left) or as list of top ranked terms (right). (G) Relative protein abundance in mESCs treated with EtOH control and 250 μM CHX for 3, 6, and 9 hours for the three most highly ranked regulator candidates per database SON (top), SPEN (middle) and GLYR1 (bottom).

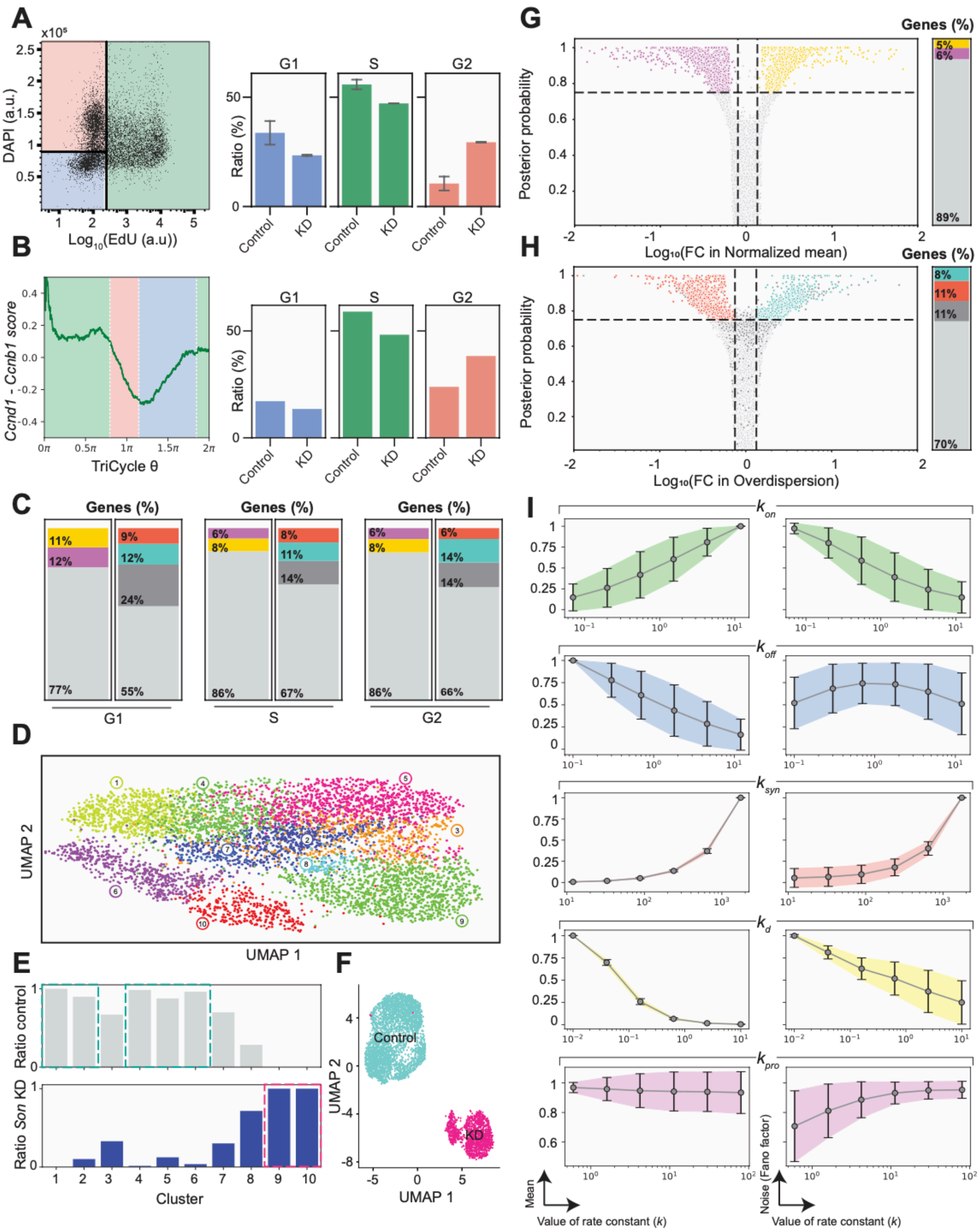


**Figure S4. Increased overdispersion upon *Son* KD is inconsistent with an extrinsic noise source, Related to Figure 3.**

**(A)** Representative scatterplot of EdU incorporation and DAPI staining, utilized to quantify the distribution of cell cycle phases in populations of cells (left). Quantification of ratio of cells in phase G1, S or G2 in control and *Son* KD, in two replicates (right).

**(B)** Relation of TriCycle θ and average ratio of single-cell z-scores of *Ccnd1* and *Ccnb1* genes. Colored areas represent the ranges of the TriCycle θ space attributed to each phase of the cell cycle (left). Ratio of cells attributed to G1, S or G2 phase from scRNA-seq data in control and *Son* KD.

**(C)** Differential expression analysis with BASiCs of the *Son* KD sample compared to the control, as in Figure 3D-E, for cells attributed to cell cycle phases G1 (left), S (center) or G2 (right). Stacked bar plot represents the % genes that display significant changes in mean (left bar) or noise (right bar).
**(D)** Clustering of all sequenced cells in both samples, control and *Son* KD, with RaceID.
**(E)** Quantification of the origin of the cells (Control or *Son* KD) in clusters defined in (D). Clusters 1,2,4,5 and 6 (highlighted in turquoise) show high proportion of cells from control sample and are defined as filtered control sample for the analysis depicted in this figure. Clusters 9 and 10 (highlighted in pink) show high proportion of cells from *Son* KD sample and are defined as filtered Son KD sample for the analysis depicted in this figure. Clusters 3, 7 and 8 present mixed populations that indicated potential failed KD in the cells belonging to the *Son* KD sample, as they cluster closely to control cells (*KD-escaping cells*). In this clusters, cells belonging to the control sample are kept while cells belonging to the *Son* KD sample are excluded.
**(F)** UMAP dimensionality reduction of sequenced cells after *Son* KD (pink) and negative control (turquoise), after filtering of *KD-escaping* cells.
**(G-H)** Differential expression analysis, for differential mean (G) and overdispersion (H) with BASiCs of filtered *Son* KD cells versus control. Volcano plot represents the fold change versus the certainty (Posterior probability) that this change is significant (significance thresholds are depicted with dashed lines). Stacked bar plot represents the % genes that display significant changes (1071 mean enhanced; 863 mean repressed; 1932 noise enhanced; 1350 noise repressed).
**(I)** Reanalysis of previously published stochastic simulations[18] of two-state telegraph model including an mRNA processing step. Simulations were performed by changing only one rate constant ($k_{on}$, $k_{off}$, $k_{syn}$, $k_d$, $k_{pro}$) at a time. Results display the mean (left) and noise (right) changes for each parameter range. Values of mean and noise are normalized to the maximum value per single parameter range and error bars represent standard deviation across the entire parameter space.

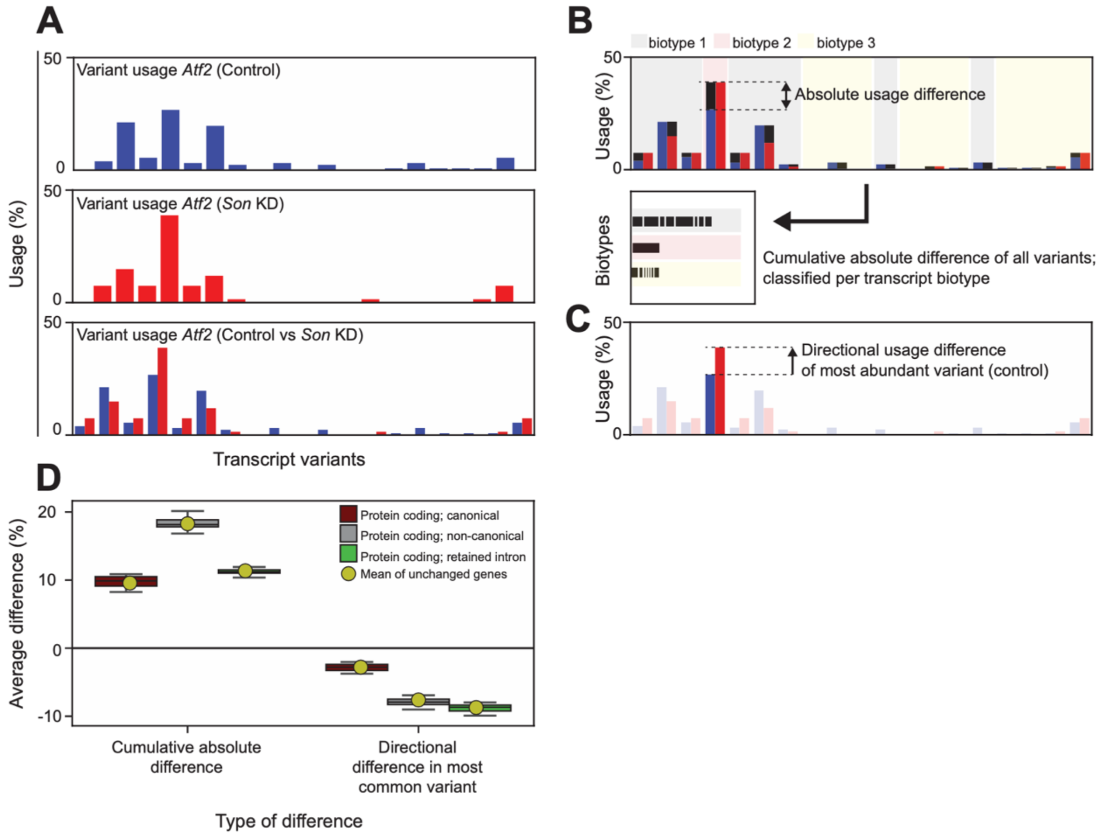


**Figure S5. Detailed analysis of isoform usage changes upon *Son* KD, Related to Figure 4.**
**(A-C)** Analysis of isoform usage difference. (A) Distribution of isoform usage for an arbitrary gene (*Atf2*) with multiple transcript variants detected in the control (top), the *Son* KD sample (middle), and a comparison between both (bottom). (B) Example of quantification of cumulative directional differences. For each transcript variant, the absolute difference in usage is calculated (top) and these absolute differences are added up depending on the biotype of each variant (bottom). (C) Example of quantification of directional difference in usage for the most common variant in the control sample. In this case, the sign of the difference indicates an increase or a decrease in usage with respect to the rest of the variants.
**(D)** Analysis of average usage difference quantified as cumulative absolute difference and directional difference in the most common variant, in 20 randomized groups of 500 genes from the entire set, depending on the biotype of transcripts (maroon, grey or green), and average difference in unchanged genes for the corresponding analysis (yellow dot).

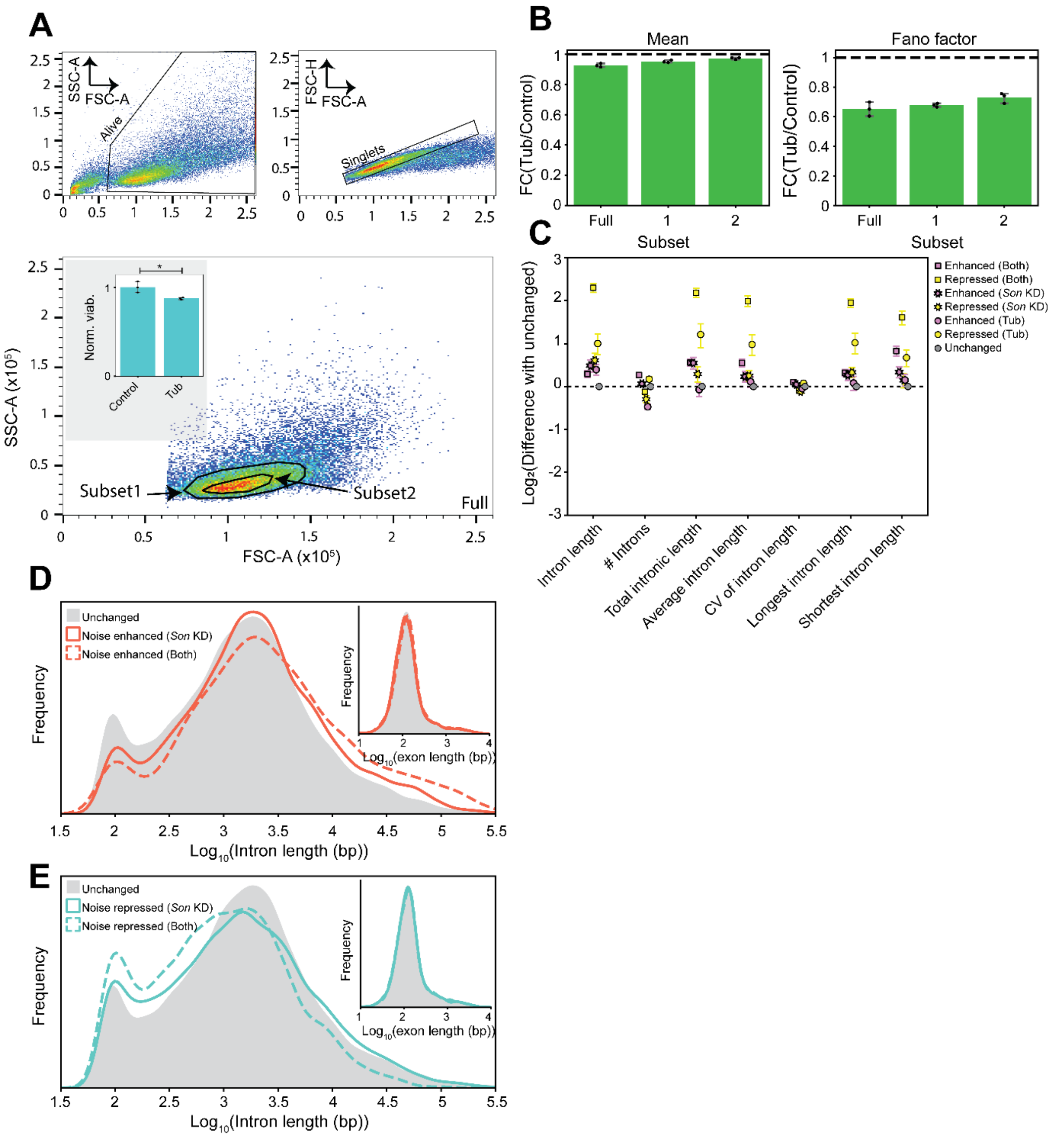


**Figure S6. Extended analysis of Tubercidin, Related to Figure 6.**

**(A)** Extrinsic noise filtering for NANOG-GFP flow cytometry upon treatment. Top left: FSC-A/SSC-A scatter plot of all events with gate for alive cells. Top right: FSC-A/FSC-H scatter plot of previously gated events with gate for single cells. Bottom: FSC-A/SSC-S scatter plot of previously gated events (full) with 2 gates for extrinsic noise filtering (Subset1 and Subset2). Bottom inset: Viability of analyzed populations as a fraction of events in the gate for live cells, normalized to the respective control sample.

**(B)** Left: Fold change in mean for one of the extrinsic noise-filtered subsets (Subset2) in control and Tubercidin, representing 3 biological replicates for the same subset. Right: Fold change in Fano factor, averaged for all biological replicates, for full single-cell population and both extrinsic noise-filtered subsets. Dashed line represents Fold change = 1.

**(C)** Association of mean regulation categories, combined from *Son* KD and Tubercidin treatment, and intron characteristics of genes in each group. Marker represents the average value for 10 randomized samples of 100 genes in a given group and for the indicated characteristic, normalized to the group of unregulated genes. Error bars represent the standard error of the mean.

**(D)** Distribution of the lengths of all introns and exons (inset) for genes in the categories of noise-enhanced in *Son* KD dataset (solid line), noise-enhanced in both *Son* KD and Tubercidin datasets (dashed line) and unchanged (grey).

**(E)** Distribution of the lengths of all introns and exons (inset) for genes in the categories of noise-repressed in *Son* KD dataset (solid line), noise-repressed in both *Son* KD and Tubercidin datasets (dashed line) and unchanged (grey).

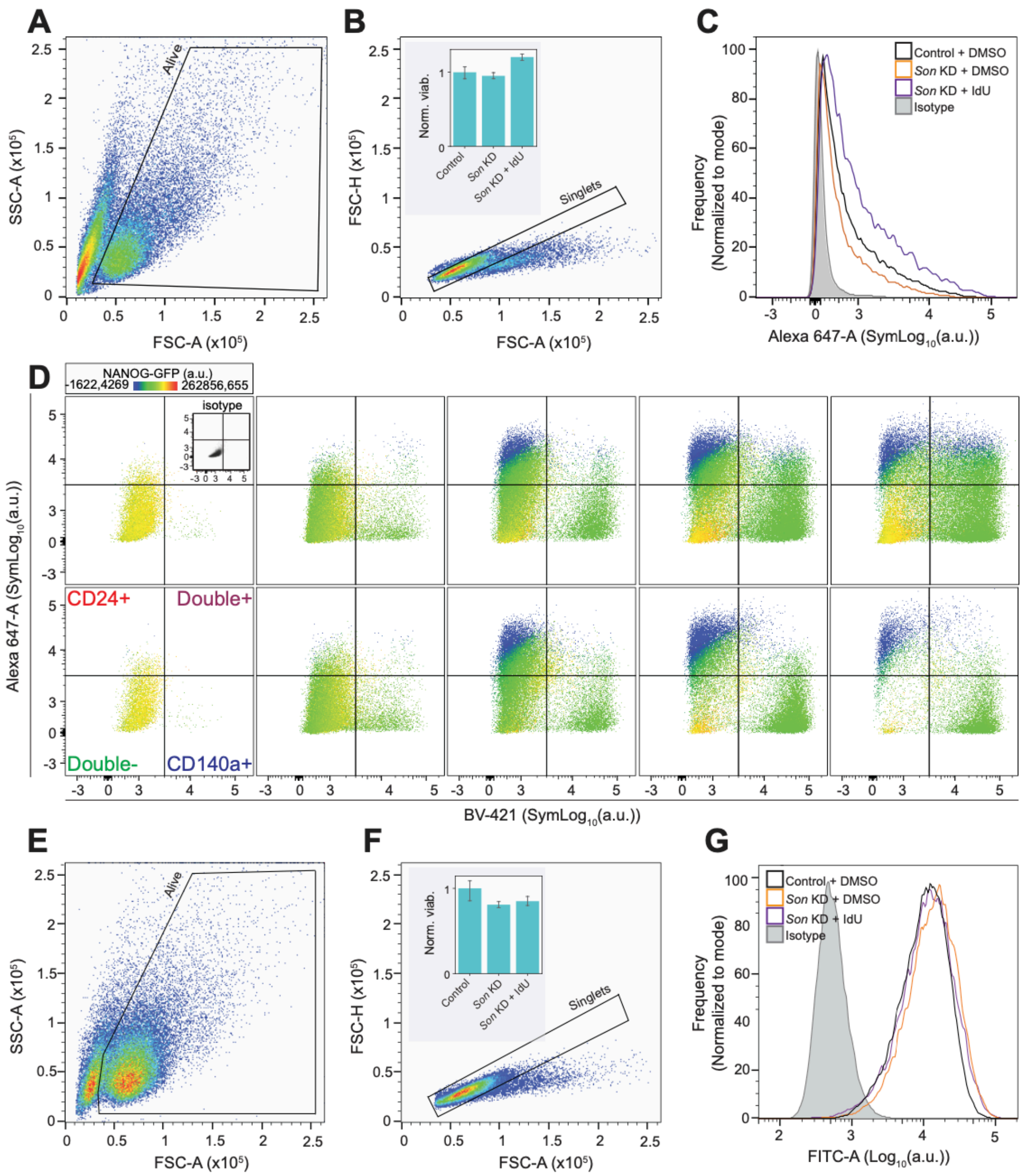


**Figure S7. *Son* KD impacts pluripotency and differentiation, Related to Figure 7.**
**(A-C)** Extended analysis of *Son* KD in CD24 expression. (A) Scatterplot of forward scatter area (FSC-A) and side scatter area (SSC-A) for a population of mESC analyzed with flow cytometry. (B) Scatterplot of FSC-A and forward scatter height (FSC-H) for the population gated in left plot. Analyzed gate selects for single cells. Insets: Viability of analyzed populations as a fraction of events in the gate for live cells, normalized to the control sample. Error bars represent 95% confidence interval. (C) Histograms of single-cell intensities of surface marker CD24 for gated population. Isotype control in grey.
**(D)** Extended analysis of how *Son* KD alters CD24, CD140a and NANOG-GFP expression at 0, 24, 48, 72 and 96 hours (left to right) of differentiation. Scatterplot of CD24 (Alexa-647) and CD140a (BV-421) expression, with NANOG-GFP expression indicated by the marker

color. Gates that define CD24+ and CD140a+ cells are defined with respective isotype controls (top left, inset).

**(E-G)** Extended analysis of how *Son* KD alters CD9 expression. (E) Scatterplot of forward scatter area (FSC-A) and side scatter area (SSC-A) for a population of mESC analyzed with flow cytometry. (F) Scatterplot of FSC-A and forward scatter height (FSC-H) for the population gated in left plot. Analyzed gate selects for single cells. Insets: Viability of analyzed populations as a fraction of events in the gate for live cells, normalized to the control sample. Error bars represent 95% confidence interval. (G) Histograms of single-cell intensities of surface marker CD9 for gated population. Isotype control in grey.
